# m6A regulates ADAR1 mediated RNA editing during macrophage activation

**DOI:** 10.1101/2025.01.28.635339

**Authors:** Valerie Griesche, Salvatore Di Giorgio, Irem Tellioglu, Chih-Yuan Kao, Julia Brettschneider, Riccardo Pecori, Johanna Schott, Georg Stoecklin, F. Nina Papavasiliou

## Abstract

Macrophages are a highly plastic innate immune cell subset that depends on environmental cues to activate, execute and resolve inflammatory responses. This plasticity of function is mirrored by the diversity of RNA modifications that dynamically decorate macrophage transcripts. Here, using the mouse macrophage line RAW 264.7 (RAW), we investigated two mRNA modifications, adenosine to inosine (A-to-I) deamination by ADAR1 and adenosine N6-methylation (m6A) by METTL3, and their interrelation.

Using short-read and single-molecule sequencing, we found that m6A deposition was unaffected by the absence of ADAR1, whereas loss of METTL3 reduced A-to-I editing levels during pro-inflammatory stimulation, suggesting functional interdependence. Co-occurrence analysis suggested that the two adenosine modifications are deposited independently on individual RNA molecules. At the same time, we find that loss of m6A methylation on the *Adar1* transcript impacts its translation and reduces ADAR1 protein levels, likely explaining the decrease in A-to-I editing in the absence of m6A. Finally, we show that loss of either METTL3 or ADAR1 impairs macrophage activation and phagocytosis in response to pro-inflammatory stimulation.

## Introduction

Dynamic mRNA modifications have a crucial impact on the fate and metabolism of a transcript, including splicing, subcellular localization, stability, and translation ^1^. N6-methyladenosine (m6A) is one of the most prominent modifications found in mRNA^1^. It is catalyzed by Methyltransferase 3 (METTL3), which forms part of the core (catalytic) m6A writer complex together with Methyltransferase 14 (METTL14) and Wilms’ tumor 1-associating protein (WTAP). The m6A writer complex has local preference for A within the consensus DRACH (D = G/A/U, R = G/A, A, C, H = A/U/C) motif ^2–5^. Within the transcript, m6A sites are enriched after the last splice site of the last exon and in the 3’ untranslated region (3’UTR) in proximity to the stop codon ^1,2,6^. Finally, m6A can also be removed from a transcript by eraser proteins fat mass and obesity-associated protein (FTO) and AlkB Homolog 5, RNA Demethylase (ALKBH5) ^7,8^. A variety of m6A reader proteins in different subcellular localizations recognize m6A and mediate m6A’s regulatory function on mRNA metabolism. Established functions of m6A include promoting transcript decay, transcript stability, translation, splicing, and nuclear export ^9–16^. Finally, recent studies note the role of m6A in preventing formation of anomalous double stranded RNA (dsRNA) structures and their recognition by innate immune sensors ^17,18^. Adenosine deamination is also a highly prominent mRNA modification. The majority of A-to-I deamination is catalyzed by ADAR1 protein which deaminates A to inosine (I) in the context of dsRNA^19,20^. Inosine pairs preferentially with cytosine and therefore is decoded as guanosine (G) by the cell’s translational machinery ^20,21^. Two different ADAR1 isoforms exist, the constitutively expressed (and nuclear) p110 isoform and the interferon-inducible (and mostly cytoplasmic) p150 isoform which plays an important role during interferon responses ^22,23^. A-to-I editing occurs most frequently in repetitive elements such as short interspersed nuclear elements (SINE), introns, and the 3’UTR, due to their propensity to form dsRNA structures^20^. Importantly, these dsRNA structures can activate cytoplasmic dsRNA sensors like MDA5 ^20^. ADAR1 binding and editing of such structures prevents activation of innate immune sensing pathways ^24,25^.

Both m6A and A-to-I deamination target adenosine, suggesting a potential for interactions. While several studies have investigated each modification independently^26–31^, questions regarding their interplay remain unresolved (and conflicting) in part because the assignment of m6A or A-to-I instances used different (and incompatible) methodologies. In our LPS/IFN-γ-stimulated RAW 264.7 macrophages, ∼60% of expressed genes (n = ∼12,800) are annotated as m6A-modified and ∼12% as harboring A-to-I editing sites, with ∼10% of genes classified as harboring both mark types (Fig. 5A). Recent advances in direct RNA sequencing have made it possible to directly assess the co-occurrence of these two adenosine modifications within individual RNA molecules. Here we revisit this problem focusing on the transcriptome of macrophages for three reasons: First, macrophages readily adapt their phenotype and function, in response to changes in tissue physiology and external stimuli. This means that we can match changes in modification events to specific (and well understood) functional outputs, from the upregulation of activation markers to the phagocytic ability of cells proficient or deficient in modification enzymes. Second, there is already a body of literature on m6A and A-to-I occurrences within macrophage transcriptomes (which can serve as a positive control, despite the fact that their interplay on individual RNA molecules cannot be assessed due to earlier identification methods). And third, macrophage cell lines (including the RAW 264.7 [RAW] cell line studied here) have been used in the past for studies on RNA processing ^32^, providing a system in which post-transcriptional regulatory mechanisms can be mechanistically assessed. Here, we generated RAW cells deficient for METTL3 or ADAR1, and used Nanopore direct RNA sequencing (validated with Illumina sequencing, where appropriate), to characterize the relationship between these two mRNA modifications and their impact on macrophage function.

## Results

### Adenosine modification enzymes differentially affect the dynamic changes that are induced within the transcriptome of RAW 264.7 macrophages under pro-inflammatory stimulation conditions

To investigate the dynamics of methylation and deamination of adenosine within the transcriptome of macrophages under pro-inflammatory stimulation conditions, we employed the murine RAW 264.7 (RAW) cell line as a model for monocyte-derived macrophages. Upon stimulation with lipopolysaccharide (LPS) and interferon (IFN)-γ, RNA was extracted at regular time intervals and examined by Illumina Next Generation Sequencing (NGS; Suppl. Fig. 1A). In line with previous observations ^33^, we detected rapid transcriptional changes characteristic of pro-inflammatory macrophage activation with upregulation of primary response genes such as pro-inflammatory cytokines Tumor necrosis factor (*Tnf*) and Interferon-β1 (*Ifnb1*) as early as 2 hours post-stimulation while secondary response and late remodeling genes such as Solute Carrier Family 7 Member 2 (*Slc7a2*), Integrin subunit alpha M (*Itgam*), or various Toll-like-receptors (*Tlrs*) 12 and 24 hours post stimulation (Suppl. Fig. 1B, Suppl. Files-Tables 1), demonstrating that the RAW cell line reacts adequately to pro-inflammatory stimuli making it a suitable model to study macrophage activation and immune responses.

To assess transcriptomic changes under these conditions, we first generated clonal cell lines lacking either ADAR1 or METTL3, by transient transfection of a plasmid coding for Cas9 and specific single guide RNAs (Fig. 1A). Wild-type (WT) cells transfected with non-targeting single guide RNAs were generated as control (termed WT throughout this study). We validated ADAR1- and METTL3-knockouts (KO) by genomic out-of-frame insertions or deletions and protein depletion via western blot (Fig. 1B-C; Suppl. Fig. 1C-H, Suppl. Fig. 2). Specifically, an antibody recognizing METTL3’s catalytic-subunit was used to confirm the absence of reported truncated (50 and 55 kDa) but catalytically active METTL3-isoforms which were not detected (Suppl. Fig. 1H) ^34^. Additionally, m6A depletion was also confirmed by Liquid Chromatography–Tandem Mass Spectrometry (LC-MS/MS) analysis of poly-adenylation (poly-A)-enriched RNA from METTL3-KO cells, confirming METTL3-KOs clones (Suppl. Fig. 1I). These experiments allowed us to generate multiple independent clones that were then further compared.

**Figure 1.**
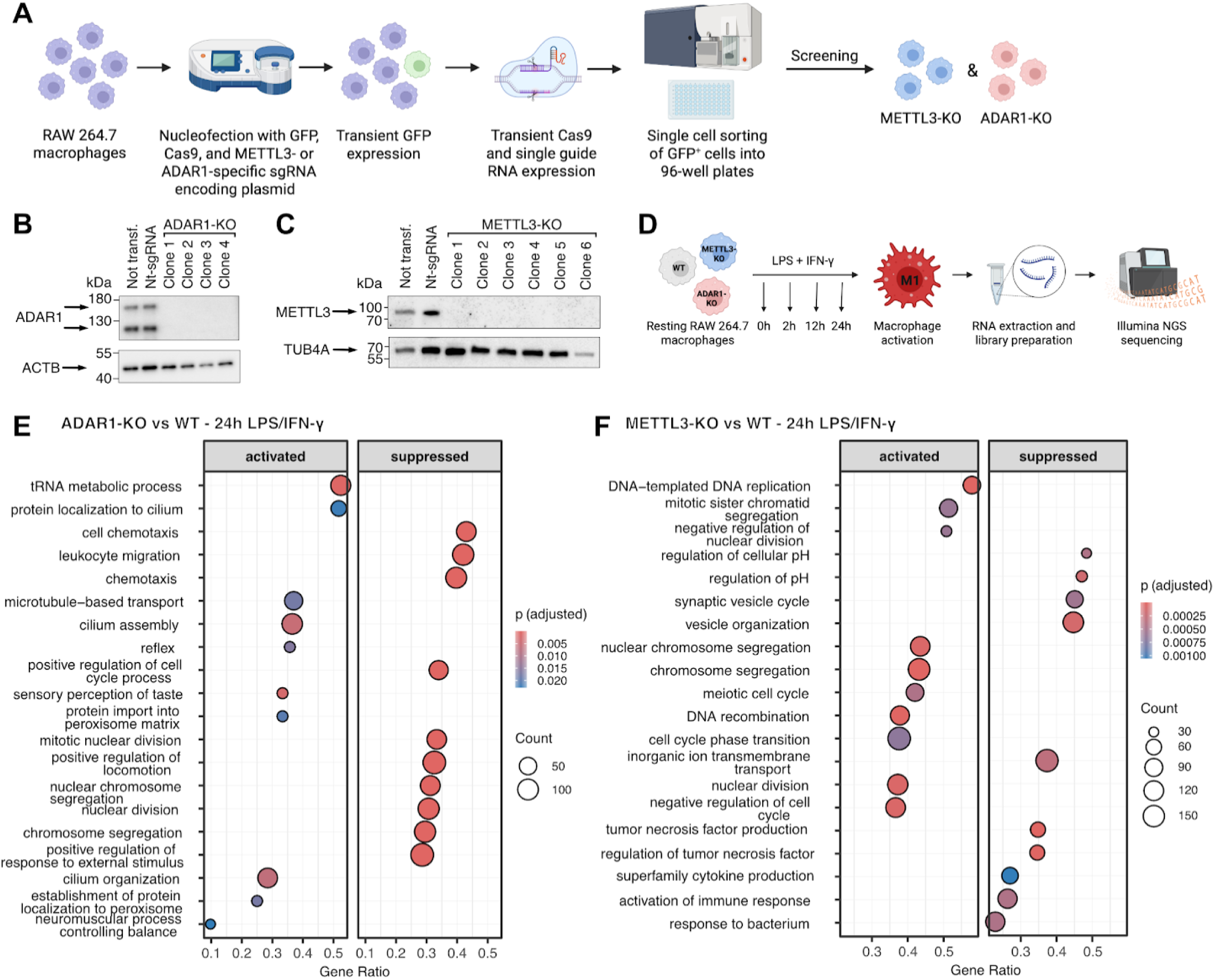
Generation and Transcriptomic Characterization of ADAR1- and METTL3-Knockout (KO) RAW Macrophage Lines. **(A)** Schematic representation of generation of clonal ADAR1-KO and METTL3-KO cell lines using plasmids encoding target specific single guide RNAs and Cas9. **(B)** Western blot validating the absence of ADAR1 protein in ADAR1-KO clonal cell lines. **(C)** Western blot validating the absence of METTL3 protein in METTL3-KO clonal cell lines. **(D)** Schematic representation of macrophage pro-inflammatory stimulation and transcripotomic analysis. **(E-F)** Top 10 most significantly enriched activated and suppressed “Gene Ontology Biological Process – (GO:BP)” terms based on Gene Set Enrichment Analysis (GSEA). The horizontal axis (Gene Ratio) shows the proportion of genes enriched in each term, while the dot size reflects the total number of genes. The color indicates the level of statistical significance (adjusted p-value). **(E)** Comparison for ADAR1-KO vs. WT at 24 h LPS/IFN-γ. **(F)** METTL3-KO vs. WT at 24 h LPS/IFN-γ. Transf, transfected; nt-sgRNA, non-targeting single guide RNA

Given that both modifications have been independently implicated in similar pathways (e.g. the prevention of aberrant dsRNA formation and innate immune activation ^17,18,24,25^), we first assessed whether the loss of ADAR1 or METTL3 also altered inflammatory pathway activity in our cell lines under resting conditions. Strikingly, the two knockouts produced divergent transcriptional responses: loss of ADAR1 was associated with suppression of an inflammatory response (Suppl. Fig. 1J; Suppl. Files-Tables 1), while loss of METTL3 led to upregulation of innate and antimicrobial immune response pathways (Suppl. Fig. 1K; Suppl. Files-Tables 1) even in the absence of external stimulation. This immune-suppressive phenotype upon ADAR1 loss contrasts with the canonical ISG upregulation described in non-immune cell types ^24,25^, but is consistent with observations in the B cell context, where reduced ADAR1 activity similarly fails to trigger the expected interferon response ^35^. In contrast, gene set enrichment analysis (GSEA) in METTL3-KO macrophages indicated upregulation of genes involved in the response to type I and type II interferons (Suppl. Fig. 1K). These findings are in line with previous studies that demonstrated interferon responses upon m6A depletion as a result from innate immune activation due to sensing of unmethylated endogenous and viral dsRNA ^17,18^.

To then study the impact of these two RNA modifications on gene expression during macrophage activation, we treated the WT and KO cell lines with LPS and IFN-γ (Fig. 1D). ADAR1-KO cells showed repressed immune functions particularly 2 hours after pro-inflammatory stimulation but also at later time points after stimulation (Fig. 1E; Suppl. Files-Tables 1), consistent with an overall suppressed immune phenotype upon loss of ADAR1 even in the absence of external stimulation (Fig. 1E; Suppl. Files-Tables 1). Conversely, in contrast to resting conditions, METTL3-KO macrophages showed impaired immune function after LPS and IFN-γ treatment. Differential expression analysis at different time points post-stimulation revealed widespread transcriptional suppression of immune-related genes in METTL3-KO versus WT macrophages, which at the pathway level translated into significant downregulation of cytokine production and immune activation programs as identified by GSEA (Fig.1F, Suppl. Files-Tables 1). Together, these findings suggest a dual role for m6A in regulating immune responses — preventing immune activation under homeostatic conditions while promoting macrophage activation in response to pro-inflammatory stimulation.

### Adenosine modification enzymes differentially affect macrophage immune function

To validate the deregulation of various pathways involved in macrophage immune function in absence of ADAR1 or METTL3, we focused on a set of genes that drive expression of immune-stimulatory cell surface markers upon pro-inflammatory stimulation, and which are known to mediate macrophage communication with other immune cells particularly T cells (Fig. 2A). Indeed, we find transcriptomic upregulation of genes such as *Cd40*, *Cd80*, beta-2-microglobulin (*B2m*, the invariant light chain of Major Histocompatibility Complex [MHC] I), MHC I (*H2-L*, *H2-K1, H2-D1*) and MHC II (*H2-Eb1*, *H2-Aa*, *H2-Ab1*) upon LPS and IFN-γ treatment in WT macrophages but also in ADAR1-KO cells with the exception of *Cd80* and *B2m* (where loss of ADAR1 resulted in a significant decrease at 24 hours LPS/IFN-γ treatment (Fig.2A). In contrast, METTL3-KO cells showed upregulation of *Cd40* and different MHC I and II chains in unstimulated METTL3-KO cells (Fig.2A), which is in line with the previously described activated baseline state upon m6A depletion (Suppl. Fig. 1K). While *Cd36* expression was downregulated in METTL3-KO cells throughout the time course, other activation markers such as MHC I chains *H2-L* and *H2-D1*, *Cd80*, *Cd40*, and *B2m* were suppressed while MHC II chains *H2-Eb1*, *H2-Aa*, and *H2-Ab1* were upregulated particularly 24 hours after LPS/IFN-γ stimulation in METTL3-KO as compared to WT macrophages (Fig. 2A). These experiments underscore the potential differential functional relevance of the two modification enzymes on macrophage function.

**Figure 2.**
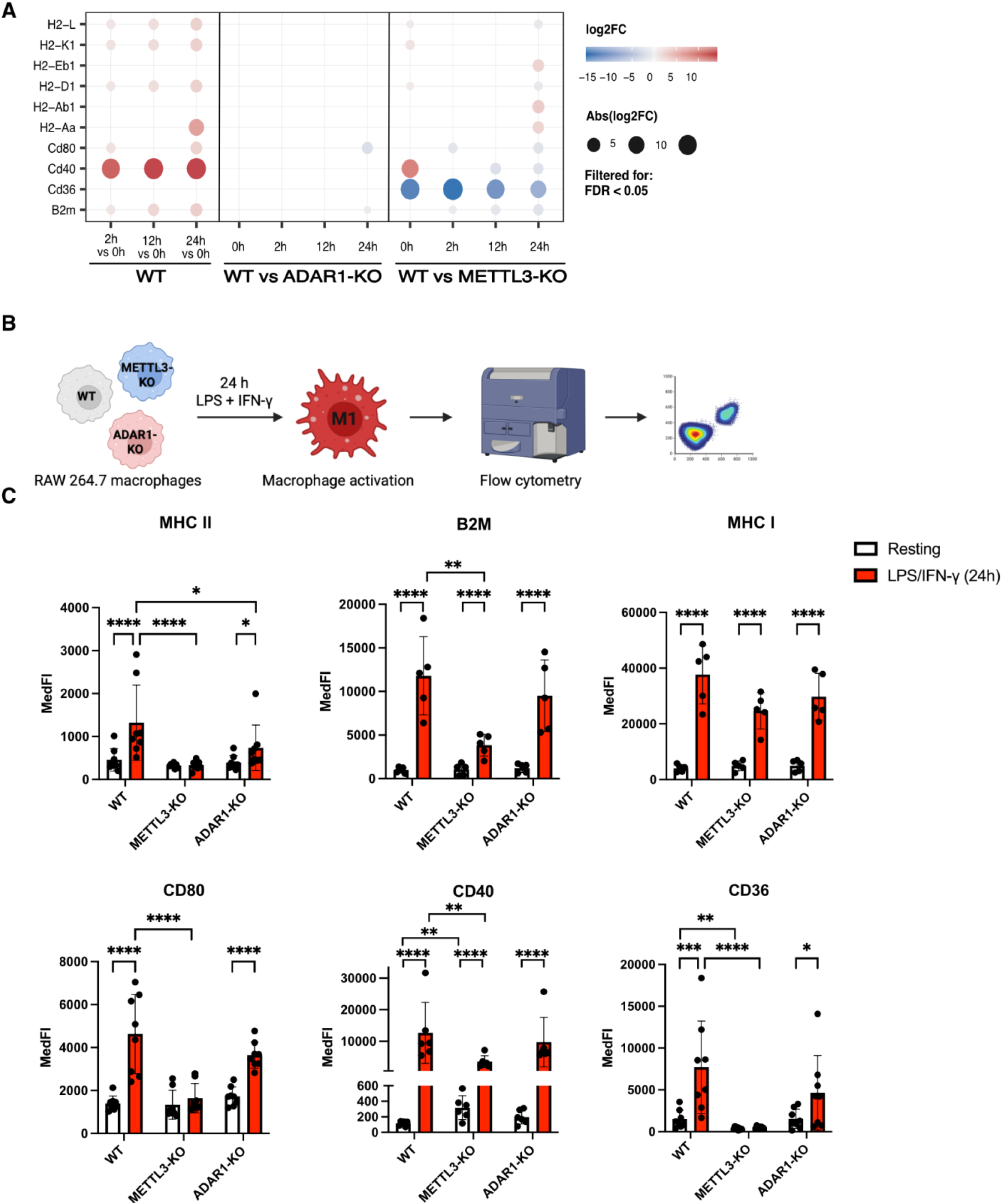
Impact of Adenosine mRNA Modifications on the Expression of Macrophage Activation Markers. **(A)** Temporal expression profiles of a subset of differentially regulated macrophage gene markers across genotypes and time points. The visualization includes genes that reached statistically significant differences in at least one comparison (FDR < 0.05). Each column represents a specific genotype (WT, METTL3-KO, ADAR1-KO) and time-point (0h, 2h, 12h, and 24h) contrast; with each genotype-time point represented by three biological replicates. The color gradient indicates the direction and magnitude of change, with red representing up-regulation and blue representing down-regulation. The size of each dot corresponds to the absolute magnitude of the log_2_ fold change, providing a clear overview of the intensity of the transcriptional response. The absence of a dot indicates that the specific gene was not significantly differentially expressed (FDR >= 0.05) for that given comparison. **(B)** Graphical representation of pro-inflammatory activation assay to assess cell surface marker expression by flow cytometry. **(C)** Cell surface expression of activation-associated markers was analyzed by flow cytometry at rest and 24h after LPS/IFN-γ treatment. Bar graphs indicate mean ± SD; CD36, CD80, MHC II: n=8; CD40: n=6; B2M and MHC I: n=5. Two-way ANOVA with Tukey’s multiple comparison test was performed on log_2_-transformed values for pairwise comparison of resting vs. stimulated within each genotype and between WT and KO genotypes at corresponding time points; only significant differences are shown. Data are displayed at the original scale (MedFI). Adjusted p-values are indicated as follows: *p < 0.05; ∗∗p < 0.01; ∗∗∗p < 0.001; ∗∗∗∗p < 0.0001. FDR = false discovery rate; SD = standard deviation; MedFI = median fluorescence intensity (y axis).

To assess whether the transcriptomic changes in macrophage activation markers were also reflected at the protein level, we measured cell surface expression by flow cytometry (Fig. 2B). Upon stimulation, WT macrophages displayed increased cell surface expression of CD36, CD40, CD80, B2M, MHC I and MHC II. ADAR1-KO macrophages showed a similar expression profile to WT macrophages with the exception of MHC II that was mildly downregulated after 24 hours LPS/IFN-γ treatment as compared to WT macrophages (Fig. 2C). In contrast, METTL3-KO macrophages showed a marked deficiency in expressing the studied surface markers as compared to WT cells upon stimulation. This differential expression was most pronounced for CD36, CD80 and MHC II, where no significant induction between resting and stimulated METTL3 -KO macrophages was observed (Fig. 2C). While most changes in gene expression were reflected at the protein level this was not always the case. For example, MHC II structural genes were upregulated in the METTL3-KO but MHC II was much reduced on the surface – possibly because loading and presentation are regulated by a broader network of genes^36^ whose dysregulation could impair MHC II surface expression. Overall, we show here that proteins like CD80 and MHC II are regulated by modification enzymes at the transcript level. These proteins play an important role in antigen presentation and co-stimulation of T cells – suggesting the functional relevance of ADAR1 and METTL3 and the modifications they “write” on macrophage mRNA.

### Adenosine mRNA modifications impact macrophage phagocytosis

As we observed pronounced changes in CD36 expression, a key scavenger receptor mediating phagocytosis, and given that global pathway analysis identified dysregulation of response to bacterium, cell killing, and vesicle organization pathways in METTL3-KO and ADAR1-KO macrophages (Fig. 1E-F, Suppl. Fig. 1J-K), we asked whether phagocytosis-related gene expression was differentially affected by loss of either modification enzyme. Phagocytosis-related genes were predominantly upregulated in WT macrophages upon LPS and IFN-γ stimulation (Fig. 3A). Loss of ADAR1 had a limited impact on this transcriptional response, with the exception of *Gas6*, which was consistently downregulated at all time points, and *Fpr2*, which was specifically reduced at 12 and 24 hours after LPS/IFN-γ treatment (Fig. 3A; Suppl. Files-Tables 1). In contrast, METTL3-KO macrophages showed broader disruption of phagocytosis-related gene expression, with specific genes up- and downregulated both at rest and upon stimulation (Fig. 3A; Suppl. Files-Tables 1). Consistent with this, GSEA using KEGG pathway annotations revealed a stimulation-dependent suppression of phagocytosis pathways in METTL3-KO macrophages (Suppl. Files-Tables 1).

**Figure 3.**
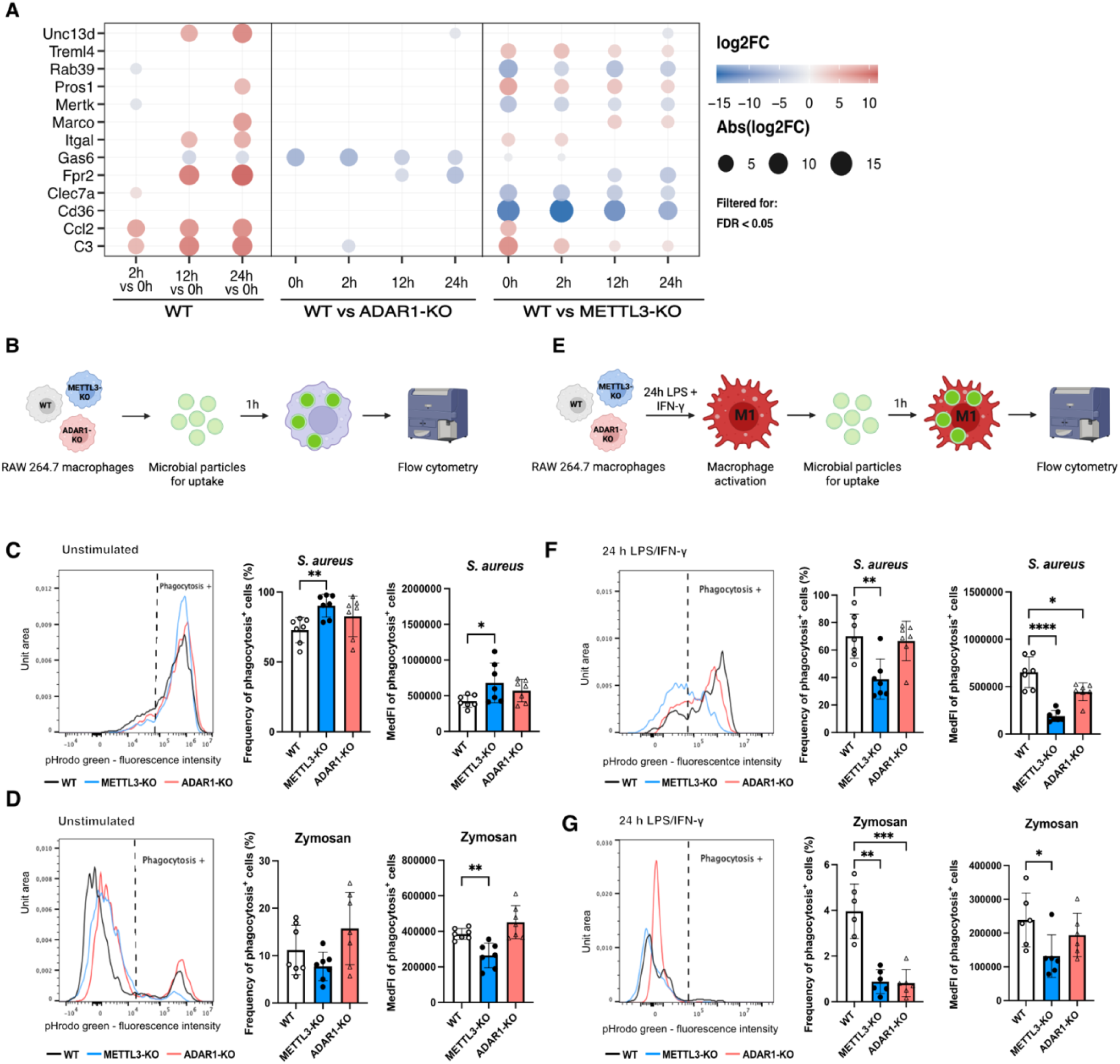
Impact of Adenosine mRNA Modifications on Phagocytosis. **(A)** Temporal expression profiles of a subset of top differentially regulated phagocytosis genes (GO:0006909:macrophages-phagocytosis) across genotypes and time points. The visualization includes genes that reached statistically significant differences (FDR < 0.05) in at least one pairwise comparison. Each column represents a specific genotype (WT, METTL3-KO, ADAR1-KO) and time-point contrast (0h, 2h, 12h, and 24h); with each genotype-time point represented by three biological replicates. Shown are the most up-regulated (top 7) and down-regulated genes (top 7) across the dataset (with some overlap). The color gradient indicates the direction and magnitude of change, with red representing up-regulation and blue representing down-regulation. The size of each dot corresponds to the absolute magnitude of the log_2_ fold change providing a clear overview of the intensity of the transcriptional response. The absence of a dot indicates that the specific gene was not significantly differentially expressed (FDR >= 0.05) for that given comparison. **(B)** Schematic representation of phagocytosis assay in macrophages without pre-treatment. **(C-D)** WT, METTL3-KO and ADAR1-KO macrophages were co-cultured with pHrodo green labeled *S. aureus* (C) and Zymosan (D) that emit fluorescent signals when they are in the acidic environment of the endophagosome. The fluorescent signal serves as a read out for phagocytic activity by flow cytometry. **(E)** Schematic representation of phagocytosis assay of macrophages pre-treated for 24h with LPS and IFN-γ. **(F-G)** WT, METTL3-KO and ADAR1-KO macrophages pre-stimulated for 24h with LPS and IFN-γ, followed by assessment of phagocytosis activity of *S. aureus* **(F)** and Zymosan **(G)** pHrodo green particles. **(C, D, F, G)** Left: exemplary histograms depicting the fluorescent signals of WT (black), METTL3-KO (blue), and ADAR1-KO (red) macrophages upon co-culture with pHrodo particles; gates for phagocytosis positive cells (indicated by dashed lines) were based on macrophage autofluorescence. Middle: Frequency of phagocytosis positive cells; one-way ANOVA with Dunnett’s multiple comparison test was performed on logit-transformed values. Right: Median fluorescence intensity (MedFI) of phagocytosis positive cells; one-way ANOVA with Dunnett’s multiple comparison test was performed on log_2_-transformed values. Bar graphs indicate mean ± SD; adjusted p-values were denoted as follows: ∗p < 0.05; ∗∗p < 0.01; ∗∗∗p < 0.001; ∗∗∗∗p < 0.0001. **(C, D, F)** n=7; **(G)** n=6.

To functionally test whether adenosine modifications impact phagocytosis, first, *resting* macrophages were incubated with two types of microbial particles conjugated with a pH-sensitive green fluorophore (pHrodo Green): *S. aureus* and zymosan (simulating yeast particles). In the acidic environment of the endophagosome, particles would emit fluorescent signals, thus allowing quantification by flow cytometry (Fig. 3B). When co - culturing unstimulated macrophages with such particles, WT and ADAR1-KO cells showed similar phagocytosis of *S. aureus* and yeast-derived zymosan (Fig. 3C-D). In contrast, METTL3-KO cells showed enhanced phagocytosis of *S. aureus* particles (Fig. 3C), possibly connected to their pre-activated state (identified by GO term enrichment; Suppl. Fig. 1K). Intriguingly, incubation of METTL3-KO cells with zymosan, led to a decrease in fluorescent signal suggesting impaired uptake (Fig. 3D). Such impaired phagocytosis of zymosan might be partially connected to reduced *Clec7a* expression in METTL3-KO cells (Fig. 3A), which is the receptor responsible for recognizing zymosan’s β-glucans and mediating uptake ^37^.

To then test if inflammatory stimulation would affect phagocytosis activity, we repeated the assay with macrophages pre-stimulated for 24h with LPS and IFN-γ (Fig. 3E). Interestingly, pre-stimulated METTL3-KO and ADAR1-KO showed significantly lower fluorescent signals in cells taking up *S. aureus* particles and the frequency of cells phagocytosing *S. aureus* was decreased in METTL3-KO as compared to WT macrophages, pointing to an impaired bacterial uptake in both KO cell lines (Fig. 3F). Similarly, the frequency of cells phagocytosing zymosan particles was strongly reduced in both KOs as compared to WT cells in LPS/IFN -γ pre-stimulated macrophages (Fig. 3G). Together, these findings indicate that phagocytosis is affected by loss of either METTL3 or ADAR1. The observed differences in uptake of *S. aureus* and zymosan between the different genotypes and inflammatory conditions likely reflect METTL3 and ADAR1’s differential regulation of distinct receptors and pathways mediating ligand recognition and phagocytosis.

### Loss of ADAR1 from RAW macrophages has negligible impact on m6A levels

After assessing the impact of modification enzymes on the function of macrophages, we turned our attention to characterizing the modifications themselves. We first profiled m6A modifications using Oxford Nanopore Technology (ONT) direct RNA sequencing under resting and stimulation conditions. Toward this, we performed differential methylation analysis in the WT (or ADAR1-KO cells) using METTL3-KO cells as background. By retaining only sites that we identified in WT but not in METTL3-KO cells, we identified METTL3-dependent sites and minimized false positives (see Methods for details on filtering criteria). Using this approach we identified a total of 89,506 m6A sites. We further validated the identified m6A sites by two orthogonal quality checks: approximately 60% of identified sites overlapped with previously reported m6A positions in the RNA Modification Database (RMBase v2.0)^38^, and 87.6% were located within the consensus DRACH motif (Suppl. Fig. 3D-F). We noted that for METTL3 transcript levels do not change with stimulation in wildtype cells, and protein levels remain largely stable (Fig. 4A-C).

**Figure 4.**
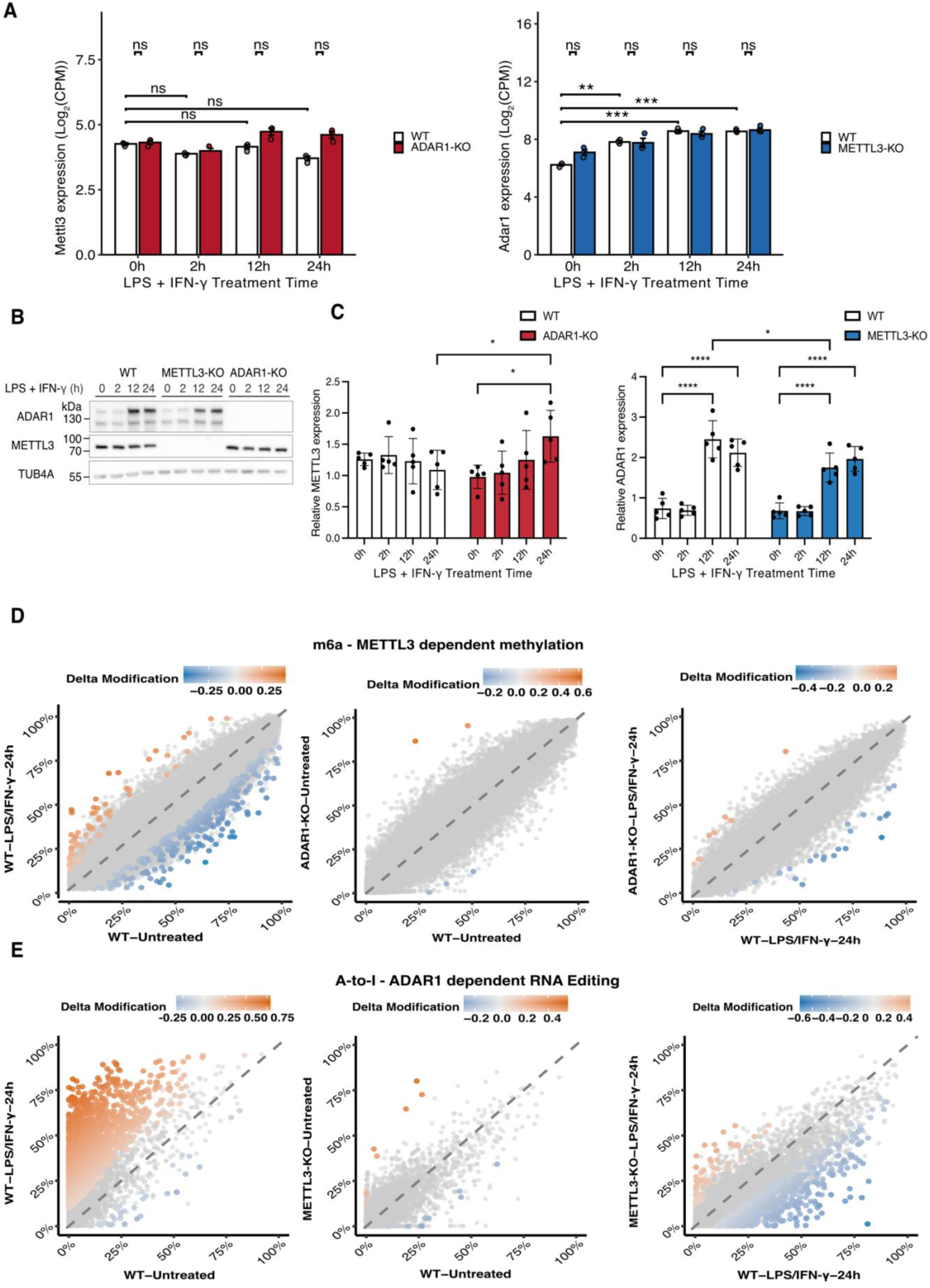
Impact of METTL3 and ADAR1 on Reciprocal Expression and Modification Dynamics. **(A)** mRNA expression of *Mettl3* (left) and *Adar1* (right) presented as the log_2_-scaled absolute expression levels of count per million CPM (CPM) across a 24-hour pro-inflammatory time course (LPS/IFN-γ, 0h, 2h, 12h, 24h). Data is shown for WT (white), ADAR1-KO (red), and METTL3-KO (blue) macrophages. Error bars represent the standard error of the mean (SEM) across biological replicates (n=3). Statistical significance was determined using edgeR differential expression analysis (see Methods). Brackets directly over individual bars indicate significant expression changes relative to the 0h baseline within that genotype, while brackets spanning genotypes indicate significant differences between KO and WT at specific time points. Significance levels: *FDR < 0.05, **FDR < 0.01, ***FDR < 0.001; ns, non-significant. **(B)** Representative Western blot showing ADAR1 and METTL3 protein expression during the time course of stimulation. **(C)** Quantification of METTL3 (left) and ADAR1 (right) protein levels presented as fold change normalized to housekeeper Tubulin Alpha 4a (TUB4A); n= 5; bars graphs indicate mean ± SD; two-way ANOVA with Tukey’s multiple comparison test was performed on log-transformed values; significant differences in pairwise comparisons between resting (0h) and individual timepoints within each genotype as well as between WT and KO genotypes at the corresponding time points are indicated by adjusted p-values as follows: *p < 0.05, ****p < 0.0001, only significant results are shown in the figure. **(D-E)** Comparison of m6A **(D)** and A-to-I editing **(E)** stoichiometry between experimental conditions. Scatterplots illustrating the absolute methylation/editing levels (mean stoichiometry, n=3) for individual sites in Group 2 (x-axis) versus Group 1 (y-axis). **(D)** Methylation: WT LPS-IFN-γ 24h vs. WT Untreated (Left), ADAR1-KO Untreated vs. WT Untreated (Middle), ADAR1-KO LPS-IFN-γ 24h vs. WT LPS-IFN-γ 24h (Right). **(E)** A-to-I Editing: WT LPS-IFN-γ 24h vs. WT Untreated (Left), METTL3-KO Untreated vs. WT Untreated (Middle), METTL3-KO LPS-IFN-γ 24h vs. WT LPS-IFN-γ 24h (Right). Each point represents a unique editing site. Sites identified as significantly differentially edited (FDR < 0.05 and absolute difference in stoichiometry Δ >= 10%) are highlighted in color, while non-significant sites are shown in grey. The diagonal identity line indicates sites with no change in editing levels between conditions.

Evaluating global modification burdens revealed a high degree of variation across biological replicates. While direct cross-genotype comparisons between control and ADAR1-deficient lines did not reach statistical significance at specific time points, a longitudinal analysis using an overdispersion-accounted GLM model revealed a significant decrease by ∼51% in global m6A burden within the ADAR1-deficient line at 24 hours of treatment (z = -4.085, *adjusted* p < 0.001, GLM quasi-Poisson with log link, Suppl. Files-Tables 2, Suppl. Fig. 4A). To evaluate the percentage of methylated adenosines at individual m6A-sites (i.e. the stoichiometry), we performed rigorous filtering retaining only sites with sufficient read coverage in all conditions (Suppl. Fig. 3A). Interestingly, global m6A stoichiometric distributions showed only weak, non-significant differences upon either stimulation or ADAR1 depletion (Suppl. Fig. 4B). This indicates that the altered global site count in stimulated ADAR1-KO cells mostly stems from genes that are not sufficiently well expressed in all conditions to pass our filter for a reliable estimation of methylation stoichiometry, rather than a widespread shift in methylation efficiency.

Given the established role of m6A in macrophage activation^26,27,31^, we further investigated site-specific stoichiometric changes. In line with overall stable stoichiometry, an evaluation of individual sites revealed that only 0.5% of the 88,850 m6A sites testable in WT (out of 89,506 detected in total; mapping across 9,671 genes) were differentially methylated during the inflammatory response. Specifically, we observed a decrease in methylation at 319 sites (within 283 genes) and an increase at 70 sites (within 48 genes; Fig. 4D; Suppl. Fig. 4E left). Additionally, m6A sites exhibit nearly identical methylation levels between WT and ADAR1 -KO macrophages, both before (5 sites differentially methylated) and after pro-inflammatory stimulation (30 sites differentially methylated; Fig. 4D middle - right; Suppl. Fig. 4E right). In summary, these data show that the m6A mark remains stable under pro-inflammatory conditions and is not affected by ADAR1 depletion (and loss of A-to-I deamination) (Suppl. Fig. 4B).

### Loss of METTL3 from RAW macrophages attenuates A-to-I editing upon pro-inflammatory stimulation

While m6A can best be detected by direct RNA sequencing (ONT), the standard for A-to-I editing is still Illumina sequencing. So we started by using the well-established Illumina sequencing platform to investigate ADAR1 dependent RNA editing events during macrophage activation, as well as to examine how these processes are influenced by the absence of METTL3. As the ADAR1 p150 isoform is interferon inducible ^22^, we expected increased levels of *Adar1* expression upon LPS and IFN-γ treatment. Indeed, WT and METTL3-KO cells exhibited an upregulation of *Adar1* mRNA already at 2 hours post-stimulation, which continued to increase at later time points (Fig. 4A right). No significant differences in *Adar1* mRNA expression were observed in absence of METTL3 after stimulation (Fig. 4A right). Protein levels followed a similar trend, albeit with a delay, as increases in ADAR1 protein were detectable only at 12 and 24 hours post-stimulation. Moreover, METTL3-KO macrophages showed a delay in ADAR1 protein upregulation in response to stimulation detectable as lower ADAR1 protein levels at 12 hours post-stimulation, and again similar protein level at 24 hours in METTL3-KO as compared to WT cells (Fig. 4B, C right).

In parallel, we investigated changes in A-to-I RNA editing burden across conditions. Following pro-inflammatory stimulation, the number of editing sites increased together with ADAR1 levels, rising from ∼0.5 editing sites per million mapped bases in resting WT macrophages to ∼1.5 editing sites per million mapped bases after 24 hours, when the highest editing levels were detected (Suppl. Fig.4C). At the same time, both in WT and METTL3-KO macrophages, a large proportion of editing sites exhibited an increase in stoichiometry upon 12 hours and 24 hours of stimulation (Suppl. Fig. 4D, Suppl. Files-Tables 2). Both observations highlight a clear editing response to inflammation. This inflammation-induced response is further supported by differential editing analysis across a total of ∼13,180 editing sites. Compared to resting WT macrophages, we observed increased editing in 17.8 % of sites (within 720 genes) at 12 hours and 31.2 % (within 1206 genes) at 24 hours, whereas a negligible fraction (0.3 %) exhibited decreased editing after 12 hours and 24 hours after stimulation (Suppl. Fig. 4D, Suppl. Files-Tables 2). When investigating the effect of METTL3-depletion on global editing burden, we observed a (non-significant) trend towards a reduced editing response in METTL3-KO compared to WT cells (Suppl. Fig. 4C). This observation was confirmed when we analyzed differential editing at individual sites. In unstimulated macrophages, METTL3-KO and WT cells showed very similar editing efficiencies, with only 8 sites exhibiting higher editing levels and 12 sites displaying reduced editing levels in METTL3-KO as compared to WT cells (Fig. 4E middle; Suppl. Fig. 4D; Suppl. Files-Tables 2). However, an increased number of sites were differentially edited between WT and METTL3-KO macrophages starting from 12 hours after stimulation. Namely, we observed a reduction in editing frequency in the METTL3-KO cells in 171 and 571 editing sites corresponding to 1.3% and 4.2 % of all editing sites, at 12h and 24h, respectively. This reduction affected 2.5 % (12 h) to 8.1 % (24 h) of genes that contain editing sites, as compared to only 30 and 56 sites showing more editing in METTL3-KO cells at 12 hours and 24 hours respectively (Fig. 4E right, Suppl. Fig. 4D, Suppl. Files-Tables 2). Overall, our data indicate that the loss of METTL3-dependent m6A leads to a modest reduction in A-to-I RNA editing levels, affecting a small subset of editing sites (1.3 % at 12h and 4.2 % at 24 h) after pro-inflammatory stimulation. While limited in magnitude, this effect suggests a possible functional interplay between m6A modifications and ADAR1-mediated RNA editing.

### Attenuation of A-to-I editing in the absence of METTL3 is not explained by a broad interdependence between m6A and A-to-I modifications

The observed dependence of A-to-I editing on m6A suggests the potential for interaction between m6A and A-to-I editing on the same transcript. As a first step, we characterized genes showing both modification types, without yet distinguishing whether both occur on the same individual molecule. In WT macrophages 24 hours post-stimulation (LPS and IFN-γ), a time point characterized by robust m6A deposition and high A-to-I editing levels, among the approximately 12,800 genes with detectable expression, more than half contained at least one modification site. Specifically, 2.0% of these genes were annotated with A-to-I editing sites only, 49.5% with m6A sites only, and 10.1% with both modification types (Fig. 5A). This indicates a significant overlap between genes displaying both modifications, with 1295 observed genes versus 923 genes expected by chance (p < 0.001, chi-squared test). Concerning their location, both modifications showed enrichment around the 3’UTR and transcript termination sites (TTS; Fig. 5B).

**Figure 5.**
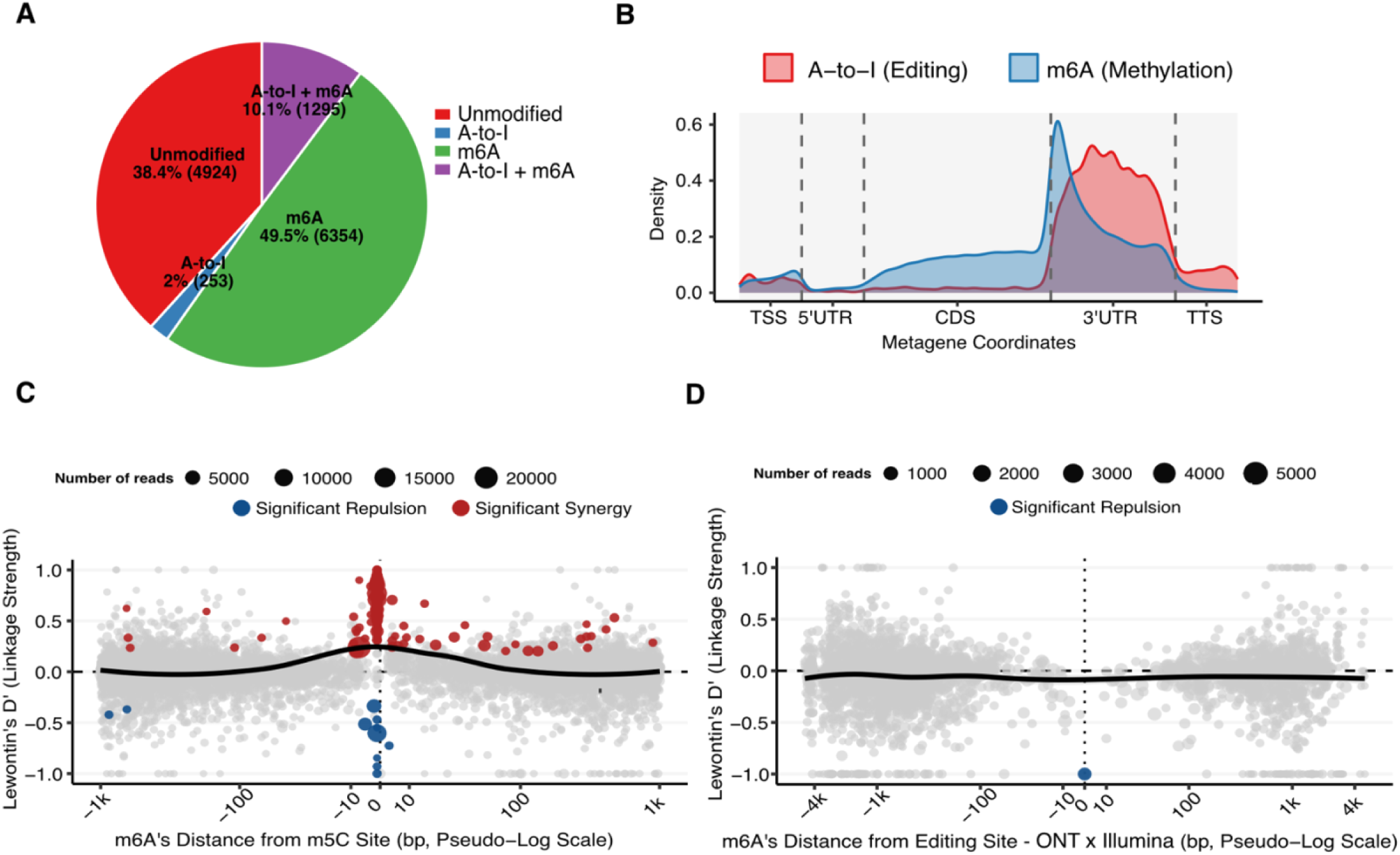
Spatial Distribution and single-molecule analysis of m6A and A-to-I Editing. **(A)** Modification status of all genes detected in the gene expression analysis (n = 12,826) in WT macrophages, combining resting and 24h LPS/IFN-γ conditions. A gene was classified as modified if it harbored at least one modification site with minimum coverage ≥10 reads and stoichiometry ≥5%, detected anywhere across the gene’s mapped reads. A-to-I editing calls were derived from Illumina sequencing; m6A calls were derived from Nanopore direct RNA sequencing. Genes are categorized as: Unmodified, A-to-I (A-to-I editing only), m6A (m6A only), or A-to-I + m6A (both modification types detected on the gene). These categories are gene-level annotation labels, not molecular descriptions: they indicate that a modification was detected somewhere within the gene’s reads, not that any single transcript or RNA molecule carries both marks, nor that all, or even most, molecules of that gene are modified. **(B)** Metagene distribution of the global localization of A-to-I editing (Illumina, red) and m6A (Nanopore, blue) sites across standardized mRNA transcript architecture. Modification site coordinates were projected onto a weighted metagene model where the coding sequence (CDS; width 3) and 3’ untranslated region (UTR; width 2) regions were scaled relative to the 5’UTR and flanking regions, transcription start site (TSS) transcription termination site (TTS), which are 1kb, to reflect biological proportions. **(C-D) Linkage and co-occurrence of RNA modifications on single RNA molecules.** Read-level linkage between proximal modification sites, quantified as Lewontin’s D′ from nanopore reads spanning both sites (per-read modification probability ≥ 0.6, ≥ 30 spanning reads). D′ = 0 indicates independence, D′ > 0 co-occurrence, D′ < 0 repulsion. Synergy and repulsion were called at FDR < 0.05 (Fisher’s exact test) and |D′| > 0.2. **(C)** m5C–m6A pairs serve as positive control to demonstrate significant non-random linkage**. (D)** m6A and A-to-I interaction was analysed using a subset of A-to-I sites that was detected in both Illumina and Nanopore sequencing (595 inosine × 1,185 m6A sites across 146 genes).

To determine whether m6A and A-to-I editing coordinate or physically interfere with one another on individual RNA molecules, we performed a linkage disequilibrium (LD) analysis by calculating Lewontin’s D’ metric ^39^. For each pair of methylation and editing sites annotated on the same gene, we identified all Nanopore reads spanning both positions simultaneously and recorded the presence or absence of each modification on every individual read. Lewontin’s D’ was then calculated as the normalized deviation between the observed co-occurrence frequency and the frequency expected under statistical independence: D’ = 0 indicates that the two modifications are deposited entirely independently on individual molecules, D’ > 0 indicates co-occurrence more frequent than expected by chance (synergy), and D’ < 0 indicates co - occurrence less frequent than expected by chance (repulsion). The magnitude of D’ therefore reflects the strength of the association and its sign the direction; significance was assessed for each pair by Fisher’s exact test with FDR correction (see Methods). This method was validated using m5C–m6A pairs as a positive control known to exhibit non-random and synergetic linkage (Fig. 5C) ^40–42^. Collectively, these single-molecule analyses demonstrate that m6A and A-to-I editing are deposited statistically independently of one another at the single-molecule level (D’ ≈ 0), neither co-occurring more often than expected by chance (synergy) nor less often than expected (repulsion), even though both marks can be found on the same molecule and can affect the same genes’ regulatory landscape (Fig. 5D; Suppl. Fig. 5A).

### Attenuation of A-to-I editing in the absence of METTL3 is consistent with a direct effect of m6A on *Adar1* transcript processing and translation

As single-molecule analysis indicated independence between m6A and A-to-I editing on individual transcripts, we examined whether the reduced A-to-I editing in stimulated METTL3-KO as compared to WT macrophages could be attributed to the direct effect of METTL3-mediated m6A deposition on the *Adar1* transcript, as reported previously ^43,44^. Analysis of our dataset confirmed the presence of several m6A sites within the last exon of *Adar1* mRNA and 3′UTR (Fig. 6A–B) that remained stable during inflammation (Suppl. Files-Tables 2) and have been proposed to contribute to the regulation of translation and ADAR1 abundance ^43^. As we observed reduced ADAR1 protein levels in METTL3-KO at 12 hours after pro-inflammatory stimulation (Fig. 4B), we utilized ribosome profiling as an orthogonal method to assess changes in translation. While WT and METTL3-KO macrophages show similar translation efficiency of *Adar1* during resting conditions, *Adar1* translation is significantly lower upon METTL3-KO as compared to WT macrophages upon pro-inflammatory stimulation (Fig. 6C; Suppl. Files-Tables 2). This suggests an inflammation-specific role of m6A in modulating *Adar1* translation efficiency during macrophage activation.

**Figure 6:**
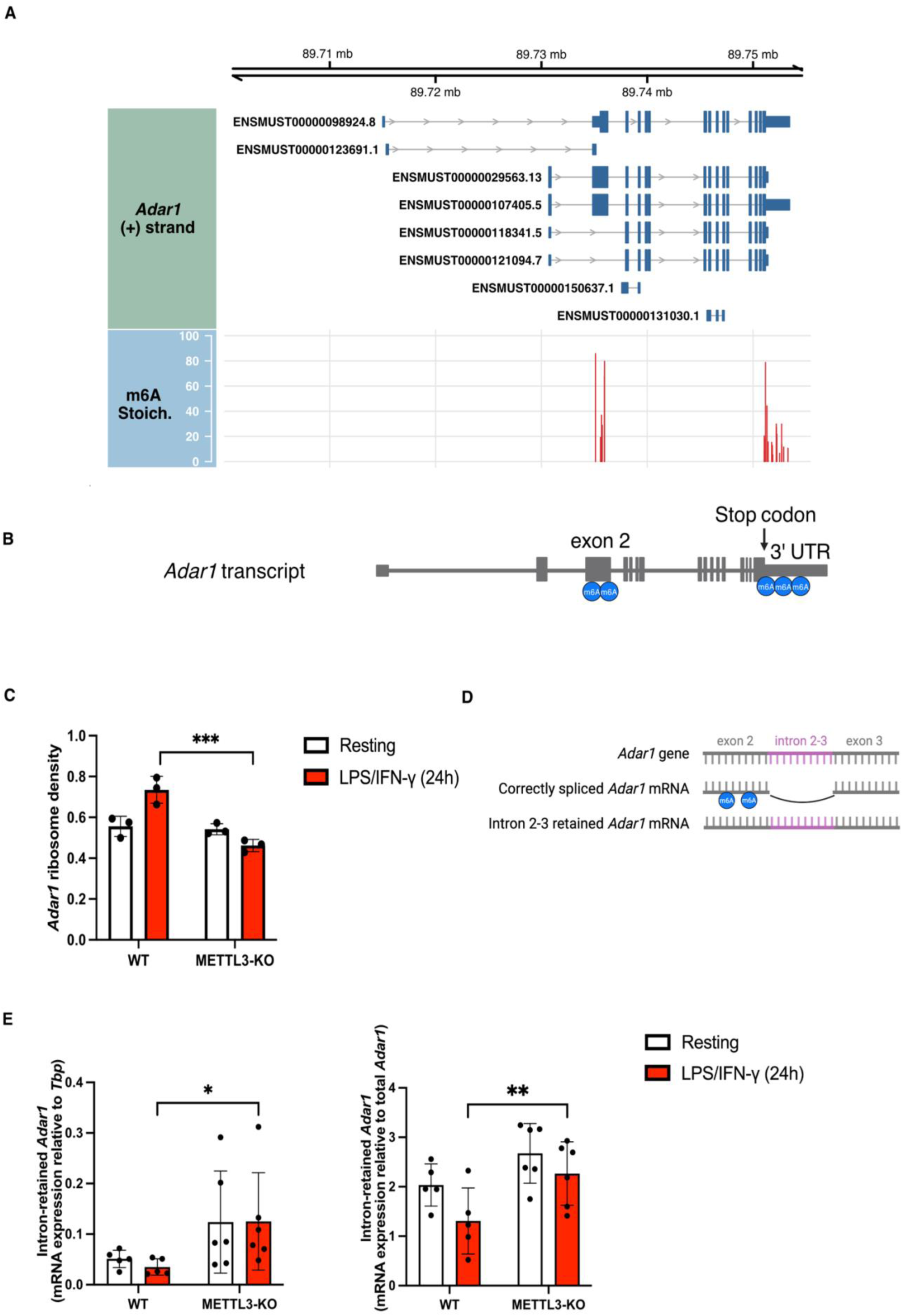
m6A Regulation of *Adar1* Transcript Processing and Translation Efficiency. **(A)** The genomic architecture and modification landscape of *Adar1* (chr3) are displayed in a strand-specific vertical stack. The top track (blue) represents the transcript isoforms on the sense strand (+), with exons shown as solid blocks. The bottom track (red) shows m6A methylation levels (%) at specific sites, revealing clusters in exon 2 and the 3’UTR. Horizontal grey grid lines denote 20% stoichiometry increments. Genomic coordinates are extended 2000 bp upstream and downstream of the gene boundaries. **(B)** Schematic representation of m6A clusters along the *Adar1* transcript. **(C)** Ribosome density of Adar1 was determined by Ribo-Seq at the resting state and 24 h after LPS + IFN-γ treatment. Values are shown as mean ± SD, n = 3. Likelihood ratio test and Benjamini–Hochberg adjustment were performed using DESeq2 **(D)** Proposed model of m6A’s role in promoting *Adar1* splicing. **(E)** Relative expression of poly-A-tailed mRNA as determined by RT-qPCR. Left: Intron 2-3 retained *Adar1* splice isoform relative to housekeeper gene TATA-box Binding Protein (TBP). Right: Intron 2-3 retained *Adar1* splice isoform relative to total *Adar1*; bar graphs indicate mean ± SD; WT: n= 5; METTL3-KO: n=6; two-way ANOVA with Tukey’s multiple comparison test was performed on ΔCt (change in cycle threshold) values; significant differences between resting (0 h) and stimulated (24 h) macrophages within each genotype as well as at the corresponding time point between genotypes are indicated. Adjusted p-values are denoted as follows: ∗p < 0.05; ∗∗p < 0.01; ∗∗∗p < 0.001. Stoich., Stoichiometry; UTR, untranslated region; RT-qPCR, reverse transcription quantitative polymerase chain reaction.

In addition to the previously reported m6A sites in proximity to the stop codon and 3’UTR, we identified a second cluster of m6A sites within the *Adar1* exon 2 (Fig. 6A-B). As m6A sites within exonic regions have been reported to affect splicing ^15,45^, we asked whether loss of m6A would affect the splicing of *Adar1* (Fig. 6D). Indeed, when analyzing mature poly-adenylated transcripts of *Adar1* by reverse transcription quantitative polymerase chain reaction (RT-qPCR), we observed increased expression levels of intron 2-3-retained *Adar1* splice isoform that was also represented by an increased ratio of intron-retained *Adar1* over total *Adar1* in stimulated METTL3-KO as compared to WT cells (Fig. 6E). As the intron-retained isoform contains premature stop codons, it would not be translated into functional ADAR1 protein, contributing to attenuated ADAR1 levels and A-to-I editing that was observed in METTL3-deficient macrophages upon LPS/IFN-γ treatment. Overall, our data suggests that the observed loss of A-to-I editing in the absence of METTL3 is due to the loss of m6A sites that decorate the *Adar1* transcript and modulate its translation.

## Discussion

Macrophages exemplify remarkable plasticity, shifting across a functional continuum to maintain tissue homeostasis and respond to diverse immunological threats. Mechanistically, plasticity requires highly orchestrated, well-regulated, and rapid responses that extend beyond direct transcriptional and translational changes, incorporating modulation of the epitranscriptome. Here we have examined the role of two epitranscriptomic modifications, m6A and A-to-I deamination, in macrophage function. Our findings reveal that the loss of m6A and A-to-I deamination leads to the deregulation of different pathways involved in inflammatory processes and immune activation (Fig. 1E-F; Suppl. Fig. 1J-K).

The importance of METTL3 and ADAR1 in regulating macrophage immune function was demonstrated by repression of transcript and protein expression of macrophage activation markers (Fig. 2) as well as impaired phagocytosis (Fig. 3) in response to LPS/IFN-γ treatment, underlining the crucial role of RNA modifications in rapid environmental sensing. Interestingly, despite the defining role of m6A in macrophage activation^26–28,31^, only 0.5% of identified m6A sites (∼88,850) showed significant stoichiometry changes upon pro-inflammatory stimulation. (Suppl. Fig. 4E). This suggests that m6A’s role in macrophage activation is not driven by changes in methylation patterns or abundance but rather, depends on the recognition of m6A marks by m6A reader proteins which influence RNA metabolism^16^ and, in turn, cellular function. Consistent with this, we and others^28^ observed significantly changed expression in many m6a readers, most prominently *Igf2bp2* in response to pro-inflammatory stimulation (Suppl. Files-Tables 1), pointing to a regulatory mechanism mediated through “m6A reading” rather than “m6A writing” dynamics.

In line with previous observations, we also found that ADAR1-mediated deamination increases after macrophage stimulation ^44,46,47^, accompanied by a global increase in the number of editing sites, which follows the rise in protein levels (in both WT and METTL3-KO cells), with minor differences observed between the two genotypes (Fig. 4B-C right; Suppl. Fig. 4B left). This results in an addition of new sites to the “editome” but also in an increase in the editing efficiency of sites that exist in resting cells (Fig. 4E left; Suppl. Fig. 4D). However this increase is clearly dampened in the absence of METTL3 (and m6A modification) suggesting an interplay between the two modifications. Such potential interplay was already detected by others in the past. For example, studies with breast cancer lines suggested that ADAR1 increased METTL3 abundance as well as transcriptome wide m6A levels, through editing a miRNA binding site in the 3’UTR of *METTL3*, thereby stabilizing its transcript ^48^. Conversely, studies in glioblastoma demonstrated that METTL3 could directly methylate *ADAR1* mRNA to promote YTHDF1-mediated translation^43^. While these and several additional studies ^26,27,29–32^ have investigated each modification independently, questions regarding their interplay had remained unresolved (and conflicting) in part because the assignment of m6A or A-to-I instances used different (and incompatible) methodologies. Recent advances in direct RNA sequencing have made it possible to directly assess the co-occurrence of these two adenosine modifications within a transcriptome (though the depth remains a technical issue that remains to be fully resolved).

Toward this, we cross-referenced m6A sites identified by Nanopore sequencing with inosine occurrences determined by Illumina short-read sequencing, finding that 10.1% of genes detectable in our gene expression dataset (out of 12,826 genes) were annotated with both modifications (Fig. 5A). Metagene analysis mapping both modifications across transcript functional units (TSS, 5’UTR, CDS, 3’UTR, TTS) revealed partial spatial overlap in the 3’UTR, despite each modification retaining a clear positional preference: m6A accumulates preferentially at the CDS–3’UTR boundary in proximity to the stop codon, while A-to-I editing displays a broader distribution extending deep into the 3’UTR, consistent with the known enrichment of SINE repeat elements in this region (Fig. 5B). While these results point to gene-level co-regulation, they cannot distinguish whether m6A and A-to-I editing act on the same individual RNA molecule, or whether they merely co-occur on the same gene without any relationship at the molecular level. To resolve this, we turned to single-molecule linkage analysis. For this we exploited the unique capacity of the Nanopore direct RNA-seq to simultaneously detect both modifications on the same read, providing a direct answer that strongly argued against obligatory co-deposition or steric exclusion at the individual molecule level, a conclusion validated by the robust positive control signal from m5C/m6A pairs (Fig. 5C). This physical independence is, in retrospect, consistent with the biochemical substrate preferences of the two enzymes, METTL3 targeting single-stranded consensus motifs and ADAR1 targeting double-stranded structure, which would not be expected to compete directly for the same adenosine in the same conformational context.

However, several important technical caveats must be acknowledged when interpreting these results. Nanopore-based Inosine detection, even with the updated base-calling model (sup.v5.2.0), remains substantially less sensitive than Illumina short-read sequencing, particularly for low-stoichiometry editing sites which are systematically missed^49^. This creates an inherent ascertainment bias in the single-molecule analysis, as only high- and medium-stoichiometry editing sites could be reliably detected. Furthermore, as visible in the single molecule analysis (Fig. 5D, Suppl. Fig. 5A), A-to-I editing sites in close proximity to m6A sites are markedly depleted in the Nanopore derived A-to-I dataset. This depletion could likely also reflect a genuine technical limitation: the Nanopore base-calling algorithm has difficulty resolving multiple distinct adenosine modifications occurring within the same short sequence window, leading to signal ambiguity and loss of modification calls precisely where the two modifications are spatially closest. This represents a critical blind spot of the current analysis and, more broadly, of current Nanopore technology. The genomic distance range where direct steric competition between m6A and A-to-I editing would be most expected is precisely the range that is least accessible to our single-molecule approach. We therefore cannot rule out that a genuine close-range antagonism exists at the molecular level, and future higher-resolution base-calling models will be essential to fully resolve the interaction landscape of modifications at short distances.

Despite this conclusion, it remained clear that loss of METTL3 also resulted in loss of A-to-I editing. We hypothesize that this is not the result of a global effect but rather of a local effect on ADAR1 translation in the absence of m6A sites that decorate the *Adar1* transcript, of which we observed several: clusters of m6A in close proximity to the stop codon and 3’UTR in the *Adar1* transcript (that were previously reported – ^43^) as well as novel sites in exon 2 which we linked to correct splicing. Both sets of m6A sites could impact translation of *Adar1*, a result which we validated both by Western (Fig.4B-C) and by ribosome profiling approaches (Fig.6C). While the stoichiometry of methylation frequency at these sites was similar during resting and during stimulated conditions, m6A’s role in promoting *Adar1* translation was only evident during pro-inflammatory conditions, coinciding with the time point when *Adar1* translation efficiency was upregulated, but not during the baseline, resting conditions. This observation is consistent with findings from Terajima et al., who similarly reported that m6A promotes *Adar1* translation efficiency selectively after IFN stimulation through YTHDF1-mediated recognition of the m6A-modified *Adar1* transcript ^44^. More generally, this indicates that stable m6A marks without changes in stoichiometry between rest and stimulation, can nonetheless produce different functional consequences for a transcript depending on the inflammatory context, possibly through stimulus-dependent changes in m6A reader expression throughout the inflammatory response.

Taken together, our results indicate that m6A and A-to-I editing do not directly compete for the same RNA molecules, within the limits of current detection sensitivity, but are nonetheless functionally coupled: m6A shapes ADAR1 abundance and, in turn, the extent of A-to-I editing during macrophage activation.

## Materials and Methods

### Cell culture

RAW 264.7 (RAW) macrophages were cultured in cell-culture treated vessels in DMEM (high glucose) supplemented with 10 % FCS, 1 % L-glutamine, and 1 % penicillin/streptomycin at 37 °C, 5 % CO2, and relative humidity 90 %. For pro-inflammatory stimulation, cells were treated with 100 ng/ml LPS and 20 ng/ml IFN-γ for indicated durations. Additionally, the purity of both cell lines was validated using the Multiplex cell Contamination Test by Multiplexion (Heidelberg, Germany). No Mycoplasma, SMRV, or interspecies contamination was detected.

### Generation and validation of RAW 264.7 knockout and non-targeting-single guide transfected cell lines

Guides targeting the genomic sequence of *Mettl3* (exon 1: GCGAGAGATTGCAGCGGCGA, exon 3: GGGCGGCAAATTTCTGGAGA), and *Adar1* (exon 2: ACTCTAACAACCCGCTGACA), as well as non-targeting guides (GCTTTCACGGAGGTTCGACG, ATGTTGCAGTTCGGCTCGAT) were cloned into plasmid pSpCas9(BB)-2A-GFP (PX458, Addgene, #48138, a gift from Feng Zhang). RAW cells were nucleofected with plasmids using Amaxa’s Cell Line Nucleofector Kit V (Lonza, 10507935) and the manufacturer’s protocol for Nucleofector II Device. 48 h after nucleofection, cells were resuspended in cell sorting buffer (PBS supplemented with 2 % FCS, 25 mM HEPES, and 2 mM EDTA), stained with Propidium iodide (Invitrogen, BMS500PI) for live/dead discrimination and GFP expressing clones were single-cell sorted for ADAR1-KO and METTL3-KO, and sorted in bulk for cells transfected with plasmids encoding non-targeting single guide (sg) RNAs (referred to as WT throughout this study) using the Aria 3 Cell Sorter (BD). After expansion, KO clonal cell lines were screened for biallelic insertions or deletions by amplification of the targeted gene region by polymerase chain reaction (PCR) using Q5 high-fidelity DNA polymerase (NEB, M0491L) with associated reagents according to NEB’s standard protocol followed by Sanger Sequencing. Knockout cell lines were confirmed on the protein level using Western blot. Unless indicated differently, each ADAR1-KO and METTL3-KO replicate presented in the experiments is derived from a single clonal cell line and each experiment contains replicates from at least 3 different clonal ADAR1-KO and METTL3-KO clones.

### Flow cytometric analysis of cell surface markers

Unstimulated or cells stimulated for 24h with 100 ng/ml LPS and 20 µg/ml IFN-γ were harvested in FACS buffer (PBS supplemented with 2 % FCS), incubated for 10 minutes in homemade Fc blocking buffer, stained for 30 minutes with antibody staining cocktail in FACS buffer, followed by 30 min staining with violet live/dead fixable staining solution (Invitrogen, 34964, 1:1000 in PBS), and fixation for 10 minutes with 4 % Paraformaldehyde (Thermo, 28908) in PBS. Between steps, cells were washed by adding an excess of FACS buffer and centrifugation for 5 minutes at 800 g at 4°C. Samples were analyzed using the Canto II (BD) flow cytometer. Antibodies used: CD40 APC/fire, Biolegend, 124632; CD36 APC, Biolegend, 102611; I-A/I-E FITC, Biolegend, 107605; anti-mouse β2-microglobulin APC, Biolegend, 154505; H-2 PE, Biolegend, 125505; CD80 PE, BD, 553769).

### Phagocytosis assay

For unstimulated samples, 100 000 cells were plated per well in flat-bottom 96-well plates in fresh medium and allowed to attach for 1 h. For pre-stimulation, 10 000 cells were plated per well in 96-well plates, and treated the next day for 24h with 100 ng/ml LPS and 20 µg/ml IFN-γ. Cells were washed twice with non-supplemented DMEM (high glucose), followed by addition of 5 µg pHrodo Green S. aureus (Invitrogen, P35367) or Zymosan (Invitrogen, P35365) particles in 50 µl non-supplemented DMEM. Cells were incubated with particles for uptake for 1 h at 37°C. Cells were harvested, stained for 30 minutes with violet live/dead fixable staining solution (1:1000 in PBS), and washed. Cells were analyzed at the Cytek Aurora Spectral Flow Cytometer using automated spectral unmixing.

### Next generation Illumina sequencing (NGS)

For NGS sequencing, RNA extraction was performed using the RNeasy Mini kit (Qiagen, 74104) according to the manufacturer. Contaminating DNA was removed using the Turbo DNase-free Kit (Invitrogen, AM1907) according to the manufacturer. RNA was quantified by Nanodrop and submitted to the DKFZ Core Facility for library preparation. rRNA was depleted and libraries were prepared using the Illumina Ribo Zero Kit (Illumina, 20037135) with 13 PCR cycles for amplification. The quality of the RNA and of the library was analysed using Agilent Tapestation 4200. Libraries were sequences for 150 cycles in paired end modus using NovaSeq 6000 on a S4 Flowcell to generate at least 60 Mio Reads per sample.

### Direct RNA sequencing using Oxford Nanopore Technology

Unstimulated and cells treated for 24h with LPS and IFN-γ were lysed in Trizol and RNA was extracted using Zymo Direct-Zol Kit (Zymo, R2050) including DNase I treatment. Libraries were prepared from 2.5 µg total RNA or from IVT RNA using the Direct RNA Sequencing Kit (SQK-RNA004) according to the manufacturer. Libraries were loaded on the PromethION RNA Flow Cell (FLO-PRO004RA) on the promethION 24 sequencing device.

### RNA LC-MS/MS

Unstimulated and cells treated for 24h with LPS and IFN-γ treatment were lysed in Trizol and RNA was extracted using Zymo Direct-Zol Kit (Zymo, R2050) including DNase I treatment. Poly-A enrichment was performed using the NEBNext Poly(A) mRNA Magnetic Isolation Module (NEB). RNA samples from three replicates were pooled and RNA was precipitated overnight at -70°C using ammonium acetate (Sigma, A2706) and ice-cold ethanol. Agilent RNA 6000 Nano Kit (Agilent, 5067-1511) was used to verify enrichment of poly-A RNA on the 2100 Bioanalyzer System.

LC-MS/MS was performed as described in detail elsewhere ^50,51^. In short, RNA was digested and an internal standard of 13C labeled yeast RNA was added. Calibration samples of unmodified and modified nucleotides were prepared. Samples were injected into the LC-MS/MS instrument for measurement. Data was analyzed using the Agilent Masshunter qualitative and quantitative software.

### Reverse transcription (RT) and RT-qPCR

RNA extraction for downstream reverse transcription was performed using the RNeasy Mini kit (Qiagen, 74104) according to the manufacturer. Contaminating DNA was removed using the Turbo DNase-free Kit (Invitrogen, AM1907) according to the manufacturer. Reverse transcription was performed using ProtoScript First Strand cDNA Synthesis Kit (NEB, E6300L) according to the manufacturer. Quantitative PCR, was performed using iTaq Universal SYBR Green Supermix (Biorad, 1725121) according to the manufacturer using the CFX Connect Real-Time System (Bio-Rad). Loading amount of samples was normalized using housekeeper TATA-binding protein (Tbp; Tbp_fw: ACCGTGAATCTTGGCTGTAAAC; Tbp_rv: TTCTTGCTGTCTTTGGAACTTTGTC) that was expressed at similar levels in all samples across conditions. Transcript expression of total *Adar1* (Adar1_ex2_fw: GATGCCCTCCTTCTACAGCC; Adar1_ex2_rv: ACTCTAACAACCCGCTGACA) and intron2-3 retained *Adar1* (Adar1_ex2_fw: GATGCCCTCCTTCTACAGCC; Adar1_in2-3_rv: TCTGGGCAGTCTCTTACCGA) were quantified.

### Western blot

Cultured cells were washed with PBS and lysed in Cell Lysis Buffer (Cell signaling, 9803) containing cOmplete, Mini, EDTA-free Protease Inhibitor Cocktail (Roche, 11836170001) and 1 mM phenylmethylsulfonylfluoride (Sigma, 93482, 1:100) for 30 minutes on ice, followed by centrifugation for 15 min at 10 000 g at 4 °C. The protein in the supernatant was quantified using the Pierce BCA Protein Assay Kit (Thermo, 23225) according to the manufacturer, and was cooked with Laemmli buffer for 5 minutes at 95 °C. Samples were loaded on Mini-PROTEAN TGX Precast Gels (4-15 %, Bio-Rad, 4561086) and run for 30 minutes at 70 V followed by 130 V until reaching desired separation. Proteins were transferred to a nitrocellulose blotting membrane (Cytiva) using BioRad wet blotting system for 70 minutes at 100 V. The nitrocellulose membrane was blocked for 1 h with 5 % skim milk in TBS-T (1X TBS with 0.1 % Tween-20). Membranes were incubated with primary antibodies (METTL3, Abcam, ab195352-1:1000; ADAR1, Santa cruz, sc-73408 - 1:500, beta-actin, Sigma, A5441 - 1:5000; alpha Tubulin, Abcam, ab4074 - 1:5000) in 5 % milk - TBS-T overnight at 4 °C while shaking. Membranes were washed three times with TBS-T and incubated with corresponding horseradish peroxidase-coupled secondary antibodies (goat anti mouse - HRP (H+L), Biorad, 170-6516; goat anti rabbit-HRP (H+L), Biorad, 170-6515; 1:5000) in 5 % skim milk - TBS-T for 1 h at room temperature while shaking. After three washes with TBS-T, membranes were incubated with chemiluminescence based detection reagent 1 and 2 at equal volumes (ECL Start Western Blotting Detection Reagent, Cytiva, RPN3243, or SuperSignal West Pico PLUS Chemiluminescent Substrate, Thermo, 34095). Chemiluminescent signals were detected using the ChemiDoc Imaging System (BioRad). Densiometric quantification was performed using the ImageJ software. The amount of protein was normalized against the amount of the housekeeper in each sample.

### Generation of *in vitro* transcribed (IVT) RNA from the poly-A transcriptome

RNA from unstimulated and 24h LPS/IFN-γ-treated macrophages was isolated in independent biological duplicates using Zymo Direct-Zol Kit (Zymo, R2050) including DNase I treatment. Poly-A enrichment was performed using the NEBNext Poly(A) mRNA Magnetic Isolation Module (NEB). For cDNA synthesis, first, 200 ng poly-A RNA, 1 µl 10 µM RT primer (5′-TTTTTTTTTTTTTTTTTTTTTTTTTTTTTTVN-3′, IDT) and 1 µl10 mM Deoxynucleotide (dNTP) Solution Mix (NEB, N0447S) were combined in 6ul and primers were annealed for 5 min at 75 °C, followed by 2 min at 42 °C. The template switching RT mix was prepared in a separate tube from 2.5 µl 4x template switching RT buffer, 1 µl 10x template switching RT enzyme mix (New England Biolabs, M0466S) and 0.5 µl 20 µM template switching oligo (5′-ACTCTAATACGACTCACTATAGGGAGAGGGCrGrG+G-3′, IDT). RNA and RT mix were combined and incubated in a thermocycler (60 min, 42 °C; 10 min, 68 °C). For Second Strand Synthesis, 50 µl Q5 Hot Start High Fidelity Master Mix (New England Biolabs, M0494S), 5 µl RNase H (New England Biolabs, M0297), 2 µl 10 µM T7 extension primer (5′-GCTCTAATACGACTCACTATAGG-3′, IDT) and 33 µl H2O were added, and incubated for 15 min at 37 °C; 1 min at 95 °C; 10 min at 65 °C). The cDNA product was cleaned using the NucleoSpin kit (Macherey Nagel, REF 740609.50) and subjected to in vitro transcription with the T7 MegaScript kit (Thermo Fisher Scientific, AM1333) according to the manufacturer’s instructions using an incubation of 4 hs at 37°C for IVT. The generated IVT RNA was cleaned up using RNA Clean and Concentrator-25 (Zymo Research, R1017).

### Ribosome profiling (Ribo-Seq)

One day before the experiment, RAW macrophages were seeded at a density of 2 × 10⁶ or 3 × 10⁶ cells per 10 cm dish for WT or METTL3-KO cells (3 different clonal METTL3-KO lines were pooled), respectively. At different timepoints after addition of 100 ng/ml LPS (Sigma, catalog no. L2630) + 20 ng/ml IFN-γ (PeproTech, catalog no. 315-05), cells were washed in ice-cold PBS supplemented with 100 µg/ml cycloheximide (Roth, Cat# 8682.3), and harvested by scraping in polysome lysis buffer (20 mM Tris-HCl buffer pH 7.4, 10 mM MgCl2, 200 mM KCl, 1 % NP-40, 100 µg/ml cycloheximide, 2 mM DTT, 1 tablet EDTA-free Roche cOmplete Mini Protease Inhibitor per 10 ml). Lysates were rotated for 10 min and then centrifuged at 9 300g for 10 min at 4 °C. About 10 % of the lysates were saved as input control. The remaining lysates were digested with RNase I (60 U per A260; Ambion, Cat# AM2294) for 20 min at 4 °C and followed 17.5–50 % sucrose density gradient centrifugation for 1 h 45 min at 40,000 rpm at 4 °C. The monosomal fractions were collected from the gradients into urea buffer (10 mM Tris-HCl pH 7.5, 350 mM NaCl, 10 mM EDTA, 1 % SDS, 7 M urea). RNA was purified by phenol extraction and precipitation in 50 % isopropanol by using phenol: chloroform: isoamyl alcohol (AppliChem, Cat# A0944) and GlycoBlue (Ambion, Cat# AM9515). rRNA was then depleted using the Human-Mouse-Rat riboPOOL Kit (siTOOLs, Cat# dp-K096-53). Input RNA was fragmented randomly by alkaline hydrolysis at pH 10.0 for 12 min at 95 °C. Both fragmented input (IN) and ribosome footprints (FP) were isolated by size-selection (25–35 nt) and extraction from a 15 % polyacrylamide Tris-borate-EDTA-urea gel. Purified RNAs were phosphorylated at their 5’ end with 10 U T4 PNK (NEB, Cat# M0201S), 40 U RNase OUT (Invitrogen, Cat# 10777019) and 1 mM ATP in T4 PNK reaction buffer for 1.5 h at 37 °C.

After end-repair, libraries were generated with the NEBNext Multiplex Small RNA Library Prep Kit (NEB, Cat# E7300) according to the manufacturer’s manual. To determine the required number of PCR cycles, 1 µl of cDNA per sample was diluted 8 times and used for a qPCR reaction (forward primer: 5’-GTTCAGAGTTCTACAGTCCGA-3’, reverse primer: 5’-CCTTGGCACCCGAGAATTCCA-3’) using SybrGreen master mix (Applied Biosystems, Cat# A25742) on a QuantStudio Real-Time PCR System. The highest threshold cycle determined among all samples was used for library preparation. The resulting libraries were purified by 10 % polyacrylamide Tris-borate-EDTA gels following the manufacturer’s instructions and sequenced on a NextSeq550 device (Illumina) with SE75 mode, acquiring an average 8 million reads per sample.

### Bioinformatic analysis of ribosome sequencing data

Sequences were demultiplexed and converted to fastq files using bcl2fastq v.2.20. Adapters were removed with the FASTX-toolkit v.0.0.13, retaining only sequences at least 20 nt long. The trimmed reads were mapped to tRNA and rRNA sequences (as downloaded from the UCSC Genome Browser) by bowtie v1.2.2, allowing a maximum of two mismatches and reporting all alignments in the best stratum (settings: -a --best --stratum-v 2). Reads that did not map to tRNA or rRNA sequences were aligned to the mouse transcriptome (Gencode VM33 as downloaded from the UCSC Genome Browser wgEncodeGencodeBasicVM33 table). Only reads between 25 and 35 nt long, and mapping to ORFs of isoforms arising from one specific gene (as defined by a common gene symbol) were counted. An offset of 12 nt upstream of the start codon and 15 nt upstream of the stop codon with respect to the 5’ end of the read was assumed. Ribosome densities for each condition were calculated from normalized read counts and compared between METTL3-KO and WT cells using DESeq2 ^52^. The downstream analysis included only those genes for which DESeq2 could calculate adjusted P values (Likelihood Ratio Test, Benjamini–Hochberg adjustment) for the fold change in ribosome density. The criteria for significantly different ribosome density was set to adjusted p <= 0.05 and absolute log_2_FC > 0.5. Data can be found in Suppl. Files-Tables 2.

### Illumina RNA-seq data processing

Adapters were trimmed using Trimmomatic v0.38 ^53^ with follow command line:

“trimmomatic.sh PE -threads 15 /${S_name}_R1.fastq.gz /${S_name}_R2.fastq.gz/Trimmed/${S_name}_R1_paired.fastq.gz /Trimmed/${S_name}_R1_unpaired.fastq.gz/Trimmed/${S_name}_R2_paired.fastq.gz /Trimmed/${S_name}_R2_unpaired.fastq.gz ILLUMINACLIP:/Illumina.fa:3:30:7 MINLEN:50”

Alignment of fastq files to the UCSC mm10 genome was performed using STAR 2 v.2.5.3.a ^54^ and the parameters detailed below.

“STAR --runMode alignReads --readFilesCommand zcat --genomeDir/UCSC_GRCm38/STAR_index_2.5.3a --readFilesIn ${S_name}R1_paired.fastq.gz ${S_name}_R2_paired.fastq.gz --outFileNamePrefix ${S_name} --runThreadN 10 -- outFilterMultimapNmax 1 --outSAMstrandField intronMotif --outSAMtype BAM SortedByCoordinate”.

Bams quality was checked using qualimap v2.2 ^55^. Output bam files were sorted and indexed using samtools v1.5 ^56^.

### Illumina RNA-editing calling and post-processing

RNA editing candidates were identified using REDItools v2.0 ^57,58^, employing the reditools.py script with the following parameters:

“reditools.py -f ${S_name}_Aligned.sortedByCoord.out.sorted.dedup.bam -o/REDITOOLED/${S_name}_Aligned.sorted_reditooled.txt -s 2 -T 2 -os 4 -m/REDITOOLED/homopol/${S_name}_Aligned.sorted_homopol.txt -c/REDITOOLED/homopol/${S_name}_Aligned.sorted_homopol.txt -r /UCSC_GRCm38/mm10.fa -sf /UCSC_GRCm38/Splicesites/mm10_splicesites.ss -q 25 -bq 35 -mbp 10 -Mbp 10”

Key parameter settings included a minimum read mapping quality of 25 (-q 25), a minimum base quality of 35 (-bq 35), and exclusion of the first and last 10 bases of each read (-mbp 10 -Mbp 10). Additional parameters specified were: strand-specific mode 2 (-s 2), strand confidence mode 2 (-T 2), and a minimum homopolymer length of 4 (-os 4).

Initial data reduction of the REDItools output was performed via a custom script (Filtering_redytool_Tables.sh). The filtration protocol exclusively retained loci characterized by an adenosine reference base and a minimum read depth of 1, thereby focusing the analysis on base-specific candidates. To establish a high-confidence baseline for non-edited sites, a custom R script (1.Make_Reditool_background.R) was developed. This script is designed to identify genomic regions that consistently exhibit no edits in the context of A-to-G transitions, representing A-to-I editing as detected by Illumina sequencing (Ctrl Positions derived from ADAR1-KOs samples). The script executes the following steps:

- Filtering: ADAR1-KO Reditool output files are processed to discard sites with more than one substitution type. Only genomic positions with a minimum coverage of 10 reads and no substitutions reported (indicated by the AllSubs column = “-”) are retained.
- Replicate and Sample Type Selection: genomic positions are required to be present in at least 2 out of 3 biological replicates and in at least 2 out of 4 sample conditions, ensuring a consistency threshold of 50 %.
- Aggregation: the script aggregates genomic positions by genomic region (Chromosome coordinates) and sample type. For each site, the median values of read coverage and the frequency of the alternative allele (expected to approximate 0) are calculated.

These steps are designed to ensure the reliability and consistency of RNA editing calls across the studied samples, enhancing subsequent filtering of false positives and SNPs.

In the next step, the in-house script “2.0.0.RNA_editing_Filtering.R” utilizes the previously generated list of genomic positions (Ctrl Positions) to identify overlapping sites in WT and METTL3-KO samples that exhibit A-to-G mutations, indicative of A-to-I editing as detected by Illumina sequencing. The filtering criteria for retaining candidate editing events involve two main stages:

#### 1. Raw Editing Sites Database Creation

This step aims at reducing the file size and optimizing subsequent steps for faster processing. The criteria applied during this stage are as follows:

- Filtering Covered Positions: genomic positions (Ctrl_Positions) are filtered to keep only those with a median coverage of ≥10 across sample replicates.
- Ctrl_Positions Matching: genomic positions are cross-referenced with the previous filtered Ctrl_Positions, and only matching sites are retained.
- Filtering Editing Candidates: Genomic positions with multiple substitution types are discarded, while positions exhibiting A-to-G substitutions and a coverage of ≥ 5 are retained.

#### 2. Refining Candidate Edited Positions

- Replicate and Sample Group Consistency: Candidate edited positions must be present in at least 2 out of 3 biological replicates and in at least 2 out of 8 sample group conditions.
- Coverage: Candidate edited positions with a median coverage of ≥10 within sample replicates are kept.
- Editing Frequency Support: Candidate edited positions are further filtered based on the median count of read supporting the editing site of ≥ 5 across sample replicates.

The script produces three output files, each tailored for specific analyses in this study: 2.Editing_sites_db_Gt10_Al5_Mc10.txt.gz: This file contains a list of edited sites filtered according to the criteria outlined above. It will be referred to as DB1 in this study.

Editing_complete_db_Gt10_Al5_Mc10.rds: This database includes a comprehensive list of genomic positions with editing observed in some samples (DB1). It highlights the presence of unedited positions overlapping with edited sites. This file is used for analyses requiring all genomic positions, such as differential editing analysis, and will be referred to as (DB2) and (DB2_an after annotation) in this study.

Editing_baseline_db_lGt10.rds: This file contains editing candidates with a coverage of ≥5. It is particularly useful for adjusting other filtering criteria, such as estimating consistency across sample groups or calculating global editing site counts and editing frequencies. This file will be referred to as DB3 in this study.

As an internal quality check, sequence motif analysis was performed on the high-confidence editing database to assess whether detected sites were enriched for the canonical ADAR1 recognition signature, characterized by guanosine depletion at the −1 position relative to the edited adenosine (see A-to-I Editing Motif Analysis). Enrichment for this motif provides supporting evidence that the identified sites are broadly consistent with genuine ADAR-mediated editing, while acknowledging that motif enrichment alone is not sufficient to exclude all false positives (Suppl. Fig. 3C).

### A-to-I Editing motif analysis

A-to-I editing sites considered for motif analysis were taken from the DB2_an, filtered to median coverage ≥10 and median editing frequency ≥5%. Flanking nucleotide sequences (±3 bp) around each validated editing site were extracted from the mouse mm10 reference genome using the BSgenome.Mmusculus.UCSC.mm10 package via the GenomicRanges^59^ and Biostrings R packages, with strand orientation handled by automatic reverse-complementation for minus-strand sites. Extracted DNA sequences were converted to RNA format (T-to-U) and assembled into a Position Frequency Matrix (PFM). Sequence logos were generated using ggseqlogo, producing both a probability logo reflecting raw nucleotide frequencies and an enrichment logo highlighting nucleotides specifically enriched or depleted relative to background (background probability = 0.25), confirming the canonical ADAR1 binding signature characterized by guanosine depletion at the −1 position (Suppl. Fig. 3C). As an additional validation, detected editing sites were intersected with the REDIportal database of known murine A-to-I editing events to quantify the proportion of novel versus previously documented sites (ref code 3.0.1Editing_Intersections_and_Motifs.R).

### Detection of differentially modified sites

Differential modification sites, specifically differential editing sites (**DES**) for Inosine (DB2_an) and differential methylation sites (**DMS**) for m6A (DB1_m6A_an), were identified using the **DSS** R package (v2)^60–62^ within our “7.0.Differential_RNA_Editing_DSS_revision.R” and “1.3.0.Differential_modification_analysis.R” codes. While originally developed for DNA methylation, DSS was implemented here using the smoothing option disabled to prevent the artificial interpolation of RNA modification signals into flanking regions. The analysis employed a rigorous Wald test based on beta-binomial distributions, allowing the test statistics to account for both biological variation (dispersion) and varying sequencing depths across samples. Candidate sites from the integrated databases (DB1m6A and DB2-Inosine) served as the primary input. For both m6A and Inosine, the log_2_ fold change (log_2_FC) was calculated to quantify the magnitude of modification frequency shifts using the following formula:

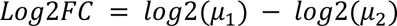

Sites were considered differentially modified at FDR < 0.05 and an absolute difference in modification frequency (|μ1 − μ2|) ≥ 0.1.

### Metagene analysis

To visualize the global distribution of m6A and A-to-I editing sites across transcript architecture, a metagene analysis was performed using a custom R script (1.5.1.Metagene_Modification_Unique_Overlaid.R). High-confidence modification coordinates from both the Nanopore m6A database and the Illumina A-to-I editing database (DB2_an and DB1_m6A_an) were converted into GRanges objects and intersected with standardized functional transcript domains, TSS flanks (-1,000 bp), 5’UTR, CDS, 3’UTR, and TTS flanks (+1,000 bp), extracted from the mouse mm10 reference transcriptome (TxDb.Mmusculus.UCSC.mm10.knownGene). To prevent artificial amplification arising from overlapping transcript isoforms, a strict deduplication step was applied, assigning each unique modification coordinate exclusively to a single functional domain. To project transcripts of heterogeneous physical lengths onto a common visual axis, modification positions within each domain were first expressed as a relative fraction (0 to 1) accounting for strand directionality (5’→3’), then projected onto a weighted pseudo-coordinate axis (range 0–8) with domain widths scaled to reflect biological proportions, the CDS assigned a width of 3 (coordinates 2–5), the 3’UTR a width of 2 (coordinates 5–7), and the remaining domains a width of 1 each. The resulting density distributions for both modifications were overlaid using ggplot2, with domain boundaries marked by vertical dashed lines and background shading, enabling direct visual comparison of the spatial preferences of m6A and A-to-I editing across the transcript body.

### Spatial Mapping and biologically-signed distance calculation

To investigate the spatial relationship and potential crosstalk between distinct RNA modifications (e.g., A-to-I editing and m6A), we calculated the biologically-signed genomic distance between paired modification events (DB4_an and DB1_m6A_an). Prior to spatial mapping, modification coordinates from both sequencing platforms (Illumina and Nanopore) were independently filtered to retain only high-confidence sites, defined by a minimum coverage of 10 reads and a modification frequency of >= 5%. Filtered coordinates were converted into GRanges objects using the GenomicRanges R package^59^. To identify proximate modifications, interval intersection was performed using the findOverlaps function, restricting the search space to a maximum genomic distance of 5,000 base pairs. To ensure that paired modifications resided on the same transcript, a strict intra-genic filter was applied, requiring perfectly matched gene identifiers, experimental conditions, and biological replicates. For all valid pairs, the absolute genomic distance was calculated by subtracting the start coordinate of the reference modification (e.g., A-to-I editing) from the target modification (e.g., m6A). Crucially, to account for transcriptional directionality, this distance was converted into a biologically-signed distance. For transcripts located on the minus strand, the absolute distance was multiplied by -1. Consequently, a negative signed distance universally indicates that the target modification is located upstream (towards the 5’ end) of the reference modification on the mature mRNA transcript. This paired coordinate database was subsequently utilized for both single-molecule phasing (Lewontin linkage disequilibrium) and continuous stoichiometric correlation analyses.

### Gene expression analysis

The quantification was performed using Salmon v0.14.2 ^63^ in mapping-based mode, providing an efficient and accurate method for transcript quantification. The following command line was used to execute Salmon:

`“salmon quant --numBootstraps 20 -i /UCSC_GRCm38/SALMON_index/ -l A -1/${Sample_name}_R1_paired.fastq.gz -2 /${Sample_name}_R2_paired.fastq.gz -p 25 - o /SALMON_expr/${Sample_name} --validateMappings”`

The Salmon outputs were imported into EdgeR^64^ via the tximport ^65^ package, which produces gene-level estimated counts and an associated edgeR offset matrix. Differential analysis for gene counts was performed using the EdgeR^64^, implemented in an in-house code. Significantly upregulated/downregulated genes were determined by abs(log_2_FC)>0.5 and FDR < 0.05; the full list of these genes and their accession numbers is provided in Suppl. Files-Tables 1. The volcano plot was generated using the EnhancedVolcano R package^66^.

Functional enrichment analysis was performed using the clusterProfiler R package^67^. Gene Set Enrichment Analysis (GSEA) was applied to the full ranked gene list, where genes were ranked by the metric sign(log_2_FC) × −log10(p-value), preserving both the direction and significance of differential expression. GSEA was performed against Gene Ontology Biological Process (GO:BP) and Cellular Component (GO:CC) terms using gene symbols, and against KEGG pathways using Entrez gene IDs, with a p-value cutoff of 0.05. Redundant GO terms were removed using the clusterProfiler simplify function. Results are displayed as split dotplots showing the top 10 activated and suppressed terms per comparison. Over-Representation Analysis (ORA) was independently performed on strictly filtered up- and downregulated gene sets (p-value < 0.05, |log_2_FC| ≥ 0.5) against GO:BP and KEGG databases, using the full set of tested genes as background universe. ORA results were visualized as dotplots, mirrored barplots, and gene-concept network (CNET) plots. (ref code: 1.0.1.Differential_Gene_expression_and_Enrichment.R)

### Nanopore direct-RNAseq pre-processing

The pod5 files were basecalled using the Oxford Nanopore Technology (ONT) basecaller Dorado v1.0.0, with the models “rna004_130bps_supv2.0@v5.2.0” for the kit “SQK-RNA004”. Basecalling was performed on a GPU (Tesla V100-SXM2-32GB).

The command line used is as follows:

“dorado basecaller --min-qscore 6 --emit-moves sup@v5.2, m5C_2OmeC, inosine_m6A_2OmeA, pseU_2OmeU, 2OmeG /ont/raw/pod5_pass/*.pod5 --reference/UCSC_GRCm38/mm10.fa --mm2-preset splice:uf:k14 > /ONT/${S_name}.bam”

The “sup, m5C_2OmeC, inosine_m6A_2OmeA, pseU_2OmeU, 2OmeG”: specifies the models used for basecalling.

The sup model (rna004_130bps_sup@v5.2.0) is the super-accuracy model for standard basecalling, while the different RNA modification models are used for detecting m6A RNA modifications.

“--mm2-preset splice:uf:k14”: configures the minimap2 alignment preset for mapping RNA reads, optimized for spliced alignment ^68^.

Output bam files were sorted and indexed using samtools v1.2.

### Nanopore modifications detection

BAM files were processed using Modkit (v0.5.0) to generate modification-specific BED files. For m6A calling in all non-IVT samples, the parameters “--filter-threshold A:0.8” and “--mod-threshold a:0.95” were applied. Additional modification thresholds were set at 0.80 for Inosine (“--mod-threshold 17596:0.80”) and 0.90 for m5C (“--mod-thresholds m:0.90”). For IVT control samples, a stringent modification probability threshold of 0.99 was maintained. These thresholds were determined empirically based on the modification probability density distributions derived from the Modkit sample-probs output (Suppl. Fig. 3B).

Modkit sample-porbs command line:

`“modkit sample-probs -t 5 --percentiles 0.0,0.25,0.50,0.75,0.80,0.85,0.9,0.95,0.99 -f 0.3 --hist --only-mapped --force --prefix ${S_name}.sorted -o ${S_name}.sorted.bam”`

Modkit pileup command line:

“modkit pileup -t 20 --log-filepath /Pileup/${S_name}.log --sampling-frac 0.4 -- max-depth 20000 --with-header --filter-threshold A:0.8 --mod-thresholds a:0.95 --mod-thresholds 17596:0.80 --mod-thresholds 69426:0.99 --mod-thresholds m:0.90 --mod-thresholds 19228:0.99 --mod-thresholds 19229:0.99 --mod-thresholds 17802:0.99 ${S_name}.bam /Pileup/${S_name}.bed”

Modkit extract calls command line:

“modkit extract calls --include-bed [file.bed] --kmer-size 9 --bgzf --force -t 6 --out-threads 2 --interval-size 10000 --queue-size 1000 --mapped-only --log-filepath [file.log] [file.bam] [file.tsv.gz]”

### ONT dRNA-seq modification preprocessing

For modification validation, we utilized respective knockout (KO) models: METTL3-KO for m6A and ADAR1-KO for A-to-I editing. In contrast, m5C analysis was conducted using IVT RNA as the negative control, a strategy necessitated by the absence of a comprehensive knockout for the methyltransferases responsible for m5C deposition in this model. Notably, IVT RNA was excluded as a negative control for m6A and Inosine calling because preliminary observations indicated that IVT samples exhibited less homogeneous transcript coverage and a distinct enrichment in 3’ UTR regions. These technical biases resulted in a lower baseline of detected modifications compared to the endogenous RNA from KO cell lines, which would have compromised the accuracy of the differential modification analysis.

Given the lack of evident divergence in modification counts between METTL3-KO and WT samples at lower probability thresholds (Suppl. Fig. 3B), a stringent post-processing framework was established to mitigate technical artifacts. The high overlap in m6A density distributions below 0.99 suggests a significant contribution of false-positive calls, potentially arising from stochastic sequencing noise or the misattribution of secondary modifications, such as m1A, to m6A signals. Similarly, for Inosine, a consistent basal signal was observed across all samples in the lower probability range.

To discover potential modification sites, we identified differentially methylated (DM) candidates under the assumption that changes in modification frequency highlight authentic sites dependent on their respective writers. We performed separate comparisons using the KO or IVT samples as negative controls:

- For m6A: We compared (1) ADAR1-KO (untreated and 24h LPS/IFN-γ) versus METTL3-KO (untreated and 24h LPS/IFN-γ), and (2) WT (untreated and 24h LPS/IFN-γ) versus METTL3-KO (untreated and 24h LPS/IFN-γ).
- For Inosine: We compared (1) METTL3-KO (untreated and 24h LPS/IFN-γ) versus ADAR1-KO (untreated and 24h LPS/IFN-γ), and (2) WT (untreated and 24h LPS/IFN-γ) versus ADAR1-KO (untreated and 24h LPS/IFN-γ).
- For m5C: High-confidence sites were identified by identifying DM candidates through the comparison of biological samples (WT) against the non-modified merged IVT samples.

These comparisons enabled the identification of potential DM sites that exhibited significant divergence between ADAR1-KO, METTL3-KO or IVT samples and the control (ctrl) across corresponding conditions.

Differential modification analysis was performed using the R package DSS (v2.54.0) within an in-house pipeline, with the smoothing option disabled. Although smoothing is optimized for CpG island analyses, where DNA methylation often exhibits strong spatial correlation, it can introduce false positives in RNA-seq data by interpolating modification signals into flanking regions where no actual methylation events were detected. For the initial DM analysis (divergent position between KOs/IVTs and WT), genomic positions were filtered based on the following criteria:

- Coverage Threshold: Only positions with a depth of ≥10 reads across all samples were retained.
- Replicate Consistency: Candidate positions were required to be present in at least 2 out of 3 biological replicates within both comparison groups.

By excluding positions showing no stoichiometry shift between target samples and the METTL3-KO (for m6A), ADAR1-KO (for Inosine) or IVT RNA (for m5C), we generated a comprehensive set of m6A, Inosine and m5C candidate sites. This dataset includes genomic coordinates, depth of coverage, and log_2_ fold-change (log_2_FC) relative to the IVT RNA or knockout samples. At this stage, no additional filters were applied based on absolute modified read counts or log_2_FC magnitude; these parameters were reserved for specific downstream analytical requirements (1.1.1.Differential_modification_analysis_baseline_file.R, 1.2.1.Differential_modification_analysis_background.R).

In the second phase, a secondary analytical bridge was implemented to transition from differential candidates to a high-confidence dataset of quantified modification levels. Using a custom R-based framework (1.2.1.Modification_Calling_using_DMA_positions_new.R), we extracted the absolute modification frequencies for all statistically validated sites (FDR < 0.05 and log_2_FC <= -0.5) across the entire experimental cohort. This directional filter specifically ensured the selection of genomic positions that were significantly less modified, or entirely unmodified, within the negative control backgrounds (METTL3-KO, ADAR1-KO, or IVT) compared to the biological samples.

Crucially, a strict biological noise filter was applied to eliminate systematic sequencing artifacts. Every candidate site was evaluated against its respective negative control; any position exhibiting a median modification frequency >= 5% in the negative control was flagged as a technical artifact and excluded from the final dataset. Retained sites were further required to show a median coverage ≥10 and a median of ≥5 modified reads in at least one biological condition. For the remaining high-confidence sites, modification percentages and coverage depths were summarized by calculating the median values across biological replicates. This produced a consolidated database (DB1m6A) containing only sites that were both statistically significant in the differential analysis and biologically suppressed in the negative control. While this multi-step pipeline drastically reduces systematic errors, the resulting output is treated as a high-confidence candidate list, acknowledging that residual artifacts inherent to direct RNA nanopore sequencing may persist.

As an internal quality check, sequence motif analysis was performed on the m6A candidate sites to assess whether detected sites were enriched for the canonical DRACH motif, characteristic of METTL3-mediated methylation (see m6A Motif Analysis) (Suppl. Fig. 3D-E). Enrichment for this motif provides supporting evidence that the identified sites are broadly consistent with genuine m6A deposition, while acknowledging that motif enrichment alone is not sufficient to exclude all residual false positives inherent to Nanopore-based modification calling.

Similarly, to confirm correct implementation of the filtering criteria requiring m6A sites to be detected across biological replicates and experimental conditions, UpSet plot intersection analysis was performed on the filtered m6A site and methylated gene databases (RMBasev2.0) (Suppl. Fig. 3F). The dominant intersection of sites shared across all conditions, comprising 52,591 m6A sites confirmed the robustness and reproducibility of the applied filtering framework.

### M6a and A-to-I editing sites effect prediction and annotation

The impact of editing sites was analyzed using the Ensembl Variant Effect Predictor (VEP)^69^, Homer^70^, and RepeatMasker^71^, integrated into an in-house R script executed via the command line (3.0.0.Comprehensive_RNA_Editing_Annotation_VEP_Homer_Repeatmasker.R). M6a sites were annotated and profiled across gene body regions using Homer and RepeatMasker. (In this study, the annotated database for editing will be denoted as DB2_an, and the annotated database for m6A will be denoted as DB1_m6A_an).

### M6a motif analysis

m6A sites considered for the motif analysis were taken from DB1_m6A (WT and ADAR1-KO samples, untreated and 24 h LPS/IFN-γ; METTL3-KO excluded), retaining sites with median coverage ≥10 and median m6A frequency ≥5% in at least one condition. Genomic sequences flanking each candidate site were extracted using the BSgenome.Mmusculus.UCSC.mm10 package, accounting for strand orientation. Sites were classified as DRACH or non-DRACH using regular expression matching — ([AGT][AG]**A**[C][ACT]) for the canonical DRACH motif. Sequence logos were generated using ggseqlogo, displaying both a standard probability logo and an enrichment logo computed by subtracting the background random probability (0.25) from observed nucleotide frequencies, thereby highlighting bases specifically enriched or depleted around m6A sites (ref code 1.2.4.Modification_Intersections_and_Motifs.R).

### Global quantification and stoichiometry comparison of RNA modification sites

For both A-to-I editing and m6A, the number of mapped bases per sample was obtained using Samtools stats and used to normalize the corresponding modification site counts, enabling reliable comparisons of global modification burdens across samples independently of sequencing depth.

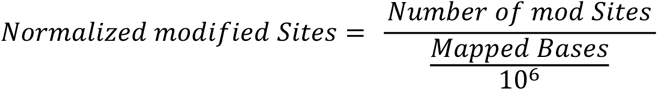

For m6A quantification, sites were counted from the high-confidence Nanopore (DB1_m6A_an) database requiring presence in at least 2 out of 3 biological replicates, a median total coverage ≥10, a median number of modified reads ≥5, and a minimum modification frequency of ≥5%. This frequency floor was specifically enforced to eliminate sequence-context-dependent Nanopore background noise; by excluding low-level technical artifacts below this threshold, significant global differences in site counts are assumed to primarily reflect true biological shifts in m6A abundance. For A-to-I editing quantification, sites were counted from DB3 applying equivalent replicate and coverage criteria (≥2/3 replicates, median coverage ≥10, median modified reads ≥5), with all positions exhibiting an editing frequency ≠ 0 included in the count. A minimum frequency floor was not applied for editing given the comparatively low background error rate of Illumina sequencing. This filtering approach nonetheless increases the potential inclusion of low-quality editing sites or false positives; however, assuming the error rate remains consistent across all samples, any observed differences should primarily reflect true biological differences in editing levels.

For both modifications, absolute site counts were modeled using a Generalized Linear Model (GLM) with a Quasi-Poisson distribution and a log link function to account for the discrete nature of count data and biological overdispersion. To correct for sequencing depth differences across samples, the natural logarithm of mapped bases was included as an offset in the model. Instead of performing all-pairwise comparisons, a set of a priori custom contrasts was defined corresponding to specific biological hypotheses. Statistical significance was assessed using General Linear Hypothesis Tests (GLHT) via the multcomp R package, with p-values adjusted for multiple comparisons using the single-step method to control the Family-Wise Error Rate (FWER), accounting for the correlation between model parameters based on the joint multivariate t-distribution.

Global stoichiometry comparisons were performed identically for both modification types. Prior to statistical modeling, sites were strictly filtered to those detected (in ≥2 of 3 biological replicates) across all sample conditions, requiring a minimum median sequencing coverage of 30, selected to mitigate the increased stoichiometry variance characteristic of lower coverage thresholds (Suppl. Fig. 3A), and a median of at least 5 modification-supporting reads, applied to both m6A and A-to-I editing sites. To analyze the stoichiometry (percentage of modified reads per genomic position) of these high-confidence sites, we employed a Beta regression model, which is highly appropriate for continuous proportional data bounded between 0 and 1. The model was fitted using a Generalized Additive Model (GAM) framework with a logit link function via the mgcv R package^72^, incorporating sample condition as a fixed effect and biological replicates as random effects. Estimated Marginal Means (EMMs) were computed on the logit scale using the emmeans R package^73^. A priori custom contrasts were tested, with p-values adjusted using the multivariate t-distribution (mvt) method to control the FWER. (ref codes: 5.0.0.Compare_n_of_editing_sites.R and 1.2.3.Modification_Stats.R).

### Co-occurrence analysis of RNA modifications – Lewontin’s normalized metric (D’)

To investigate the potential spatial coordination or interference between distinct RNA modifications on the same RNA transcript (e.g., m6A and A-to-I editing), we performed a read-level linkage disequilibrium (LD) analysis using long-read nanopore sequencing data. Modification calls and their corresponding probabilities at single-molecule resolution were extracted from modkit-processed alignments.

Prior to LD calculation, modification sites were filtered based on stoichiometry and spatial proximity. Sites were categorized into “Low,” “Medium,” or “High” modification frequencies based on the lower (0.25) and upper (0.75) quartiles of the observed modification frequencies across the dataset. Only site pairs where both modifications exhibited “Medium” or “High” frequencies were retained for pairwise analysis, ensuring that evaluated sites had sufficient modification penetrance to detect meaningful interactions.

For each pair of proximal modification sites, we extracted reads spanning both loci. Using modification genomic positions previously identified as reported in the previous sections, the modification probability per site and read level was extracted (modkit extract calls) and a probability threshold of >= 0.6 was required to call a site as modified. Furthermore, only pairs with a minimum read coverage of 30 spanning both sites were included in the statistical analysis.

Linkage disequilibrium was quantified using Lewontin’s D’, which normalizes the raw linkage disequilibrium (D) by the theoretical maximum possible LD (D_max_) given the individual modification frequencies.

Raw LD (D) was calculated as:

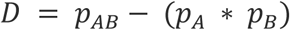

where p_A_ and p_B_ represent the overall modification frequencies at site A and site B, respectively, and p _AB_ represents the observed frequency of their co-occurrence on the same physical read. To account for the dependency of D on the individual modification frequencies, the normalized metric D’ was calculated as:

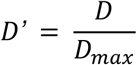

where D_max_ is defined piecewise based on whether the raw linkage (D) indicates repulsion or synergy:

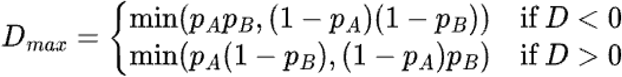

To determine the statistical significance of the observed linkage for each pair, a Fisher’s Exact Test was performed on the 2 x 2 contingency table of modification states (modified vs. unmodified for site A and site B) across all spanning reads. P-values were subsequently adjusted for multiple comparisons using the False Discovery Rate (FDR) method. Site pairs were classified as exhibiting “Significant Synergy” if they possessed an FDR < 0.05 and D’ > 0.2, and “Significant Repulsion” if they possessed an FDR < 0.05 and D’ < -0.2.

### Replicates and statistics

All replicates presented in the graphs and used for statistical analysis throughout the study are biological replicates that were generated from independent cell cultures (cultured, treated and harvested independently). For a subset of samples, the initial Nanopore sequencing runs did not yield sufficient coverage. To achieve the minimum sequencing depth (∼7M of mapped reads) required for downstream analysis, these samples were sequenced a second time on additional flow cells. The resulting reads from these technical replicates were then merged and treated as a single pooled biological replicate.

For experimental data, statistical analysis was performed using GraphPad Prism in-built analysis tool. Log-transformed relative protein expression (determined by Western blot), Δct values for mRNA expression (determined by RT-qPCR), and log_2_-transformed MedFI cell surface markers (determined by flow cytometry) were statistically tested by 2-way Anova with Tukey’s multiple comparison test. Log_2_-transformed MedFI and logit-transformed frequencies (fractions) of cells in phagocytosis assay were compared using one-way Anova with Dunnett’s multiple comparison test. For visualization, data values were displayed on the linear scale. Details on experimental values, statistical test results and significance values depicted in the corresponding figures can be found in Suppl. Files-Tables 1. Details on statistical testing in the bioinformatic data are described above in the respective subsection in the methods. Significant values are reported in the figures as adjusted p values as follows: ∗p < 0.05; ∗∗p < 0.01; ∗∗∗p < 0.001; ∗∗∗∗p < 0.0001.

### Computational tools and AI-assisted methods

R, and GraphPad Prism were used for data analysis, statistical analysis, and data visualization. Flow cytometric analysis was performed using FlowJo. Artificial intelligence–assisted tools were used to support code development, language refinement, and proofreading of the manuscript. All scientific content, analyses, interpretations, and conclusions remained the responsibility of the authors.

## Data availability

The raw sequencing data are available under BioProject accession PRJNA1232413. The code used for this manuscript is available at https://github.com/Salvobioinfo/M6a_AtoI_interplay. Additional data required to reproduce the analyses are deposited at https://doi.org/10.6019/S-BSST3186.

## Acknowledgments

We thank the Next Generation Sequencing Core Facility and the Flow Cytometry Core Facility, German Cancer Research Center (DKFZ), for providing excellent sequencing and sorting services. We thank the Helm and Hengesbach working groups, particularly Martina Schmidt-Dengler and Martin Hengesbach, for their assistance and guidance in the LC-MS/MS experiment. We gratefully acknowledge funding by: the Deutsche Forschungsgemeinschaft (DFG, German Research Foundation) – RTG2727 – 445549683 (to VG), by SVB TRR319-RMaP (FNP, VG, CK and GS); by the HI-TRON Kick-Start Seed Funding Program 2021 awarded to RP; and finally, by the Baden-Württemberg Stiftung (BWSt) [BWST-EPI-050] to S.d.G.

## Authors contribution

VG, SdG, FNP, and GS conceived the idea of the work. VG created the clonal RAW knockout cell lines and performed all experiments presented in this manuscript with the exception of sample processing and data analysis of Ribosome Sequencing performed by CK. SdG pre-processed the Nanopore Sequencing data. SdG performed all bioinformatic analyses regarding the Illumina and Nanopore Sequencing data. JB assisted and provided guidance regarding statistical analyses. RP, FNP and SdG provided scientific guidance for the manuscript. VG, SdG, and FNP wrote the manuscript, which was reviewed by RP, GS, and CK. All authors consented to the final version of the manuscript.

## Supplementary Material for

**Suppl. Fig. 1.**
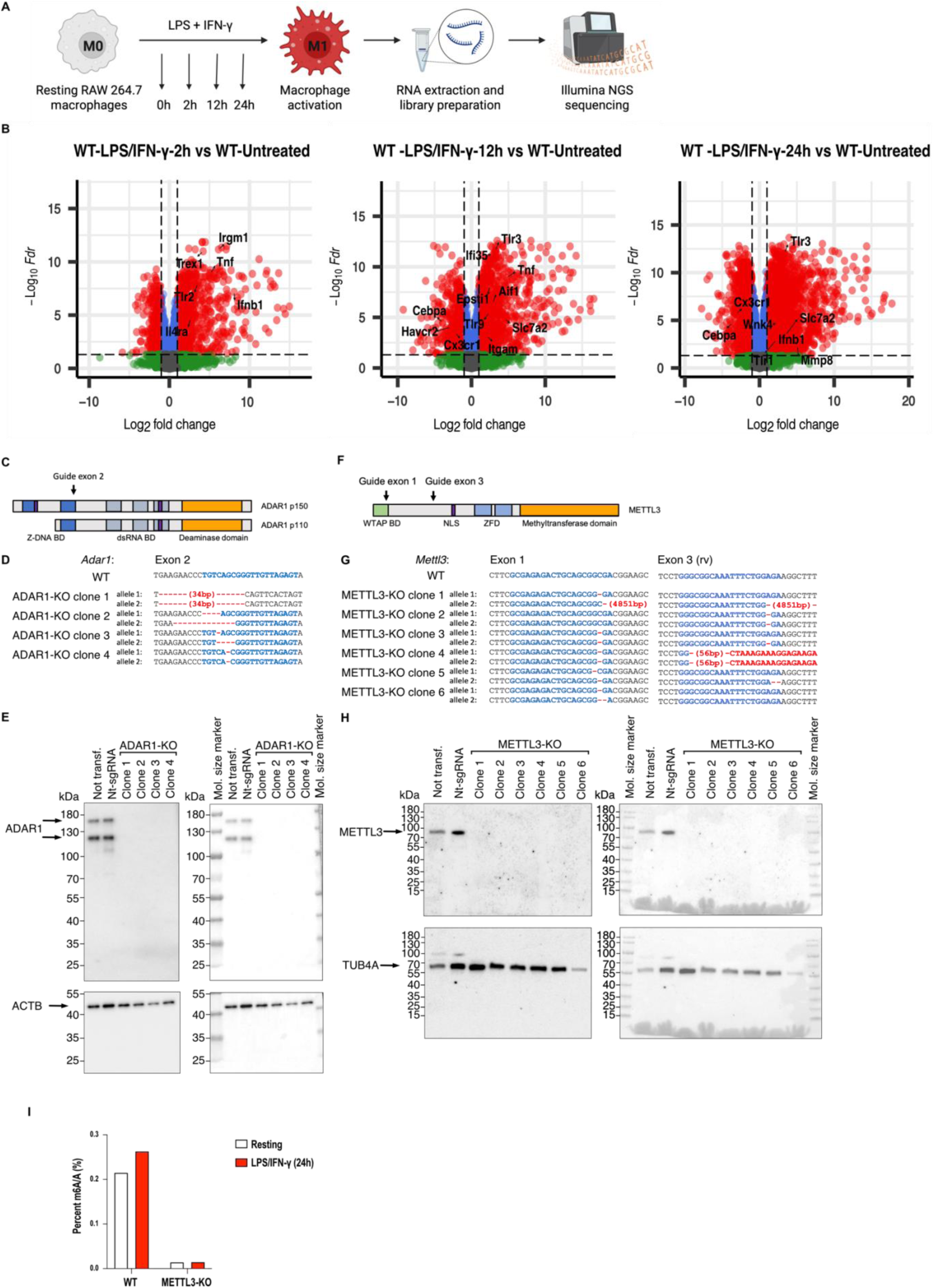
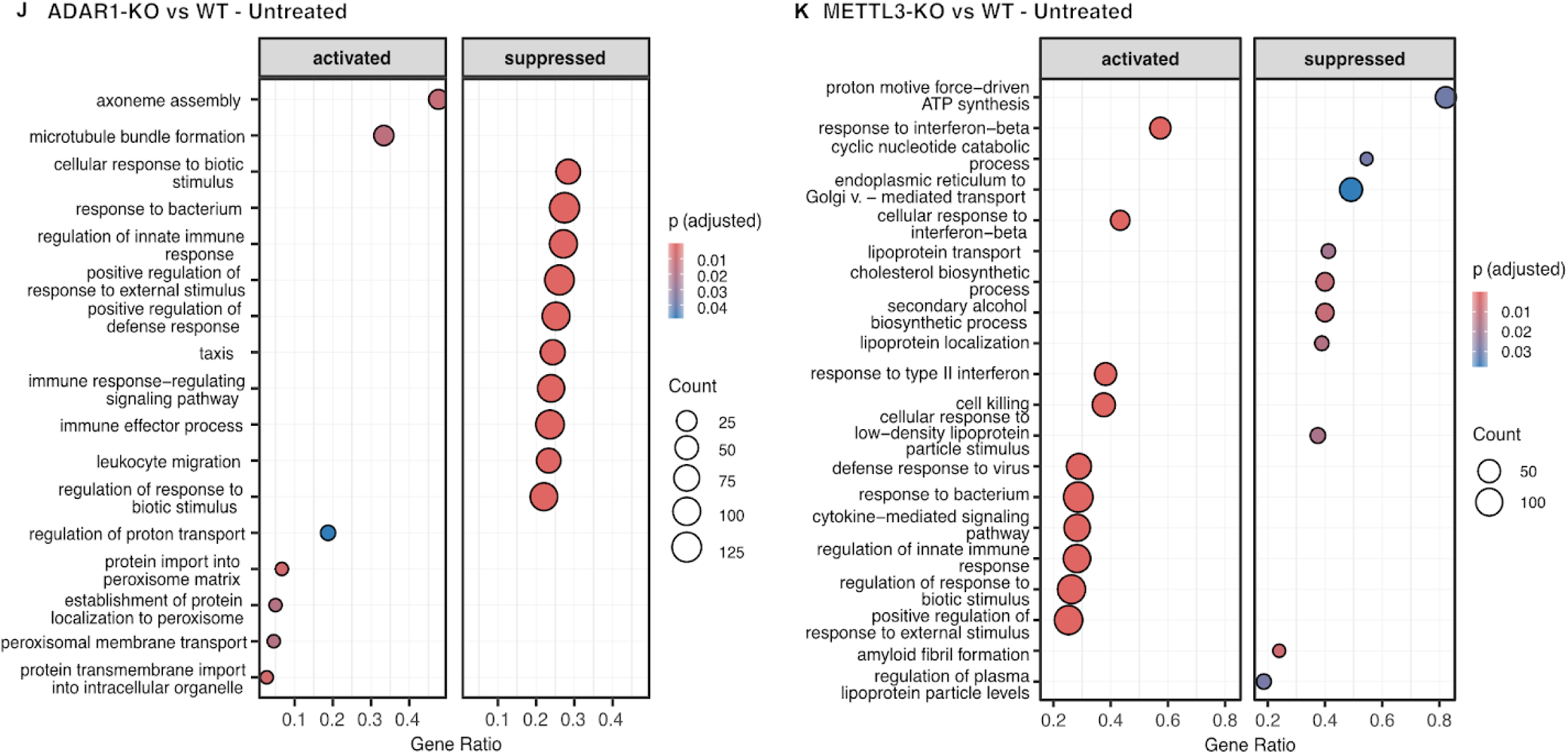
Characterization of ADAR1-KO and METTL3-KO Macrophage lines and baseline WT transcriptional response. **(A)** Schematic representation of macrophage pro-inflammatory stimulation and transcriptomic analysis. **(B)** Progression of the inflammatory response in wild type (WT) cells at 2h, 12h, and 24h post-stimulation compared to the untreated (0h) state. Volcano plots illustrating differentially expressed genes (DEGs). The x-axis indicates Log_2_FC and the y-axis represents -Log_10_(FDR). Red points denote significantly regulated genes (FDR < 0.05 and abs(Log_2_FC) > 1). Top-ranked up- and down-regulated genes are highlighted with labels; n=3. **(C)** Depiction of single guide RNA-targeted region in *Adar1*. **(D)** Clonal ADAR1-KO lines were analyzed for insertions or deletions in targeted region in *Adar1* by Sanger sequencing of PCR-amplified genomic sequences. Changes in genomic sequences are displayed; n=1. **(E)** Uncut Western blot membrane showing the absence of ADAR1 protein in ADAR1-KO lines as compared to not transfected (not transf.) and cells transfected with non-targeting single guide RNAs (nt-sgRNA); left: chemiluminescent signal; right: overlay with molecular size marker; n=1. **(F)** Depiction of single guide RNA-targeted regions in *Mettl3*. **(G)** Clonal METTL3-KO lines were analyzed for insertions or deletions in targeted region in *Mettl3* by Sanger sequencing of PCR-amplified genomic sequences. Changes in genomic sequences are displayed; n=1 **(H)** Uncut Western blot membrane showing the absence of METTL3 protein in METTL3-KO lines as compared to not transfected and cells transfected with nt-sgRNA; right: overlay with molecular size marker; n=1. **(I)** Liquid chromatography-tandem mass spectrometry of RNA enriched for polyadenylated transcripts to quantify m6A levels in WT and METTL3-KO cells; n=1. **(J-K) Basal Pathway Dysregulation in KO Lines.** Gene Set Enrichment Analysis (GSEA) of GO Biological Processes in the resting state (0h). Dot plots illustrate the top significantly enriched terms (q < 0.05) partitioned by activation status (Activated vs. Suppressed); n=3. (J) Comparison of ADAR1-KO Untreated vs. WT Untreated. (K) Comparison of METTL3-KO Untreated vs. WT Untreated. The x-axis represents the Gene Ratio (proportion of core enrichment genes within each term). Dot size is proportional to the gene count, and the color gradient represents the adjusted p- value. FDR, false discovery rate; BD: binding domain; NLS: nuclear localization signal; ZFD: zinc finger domain; mol.: molecular.

**Suppl. Fig. 2.**
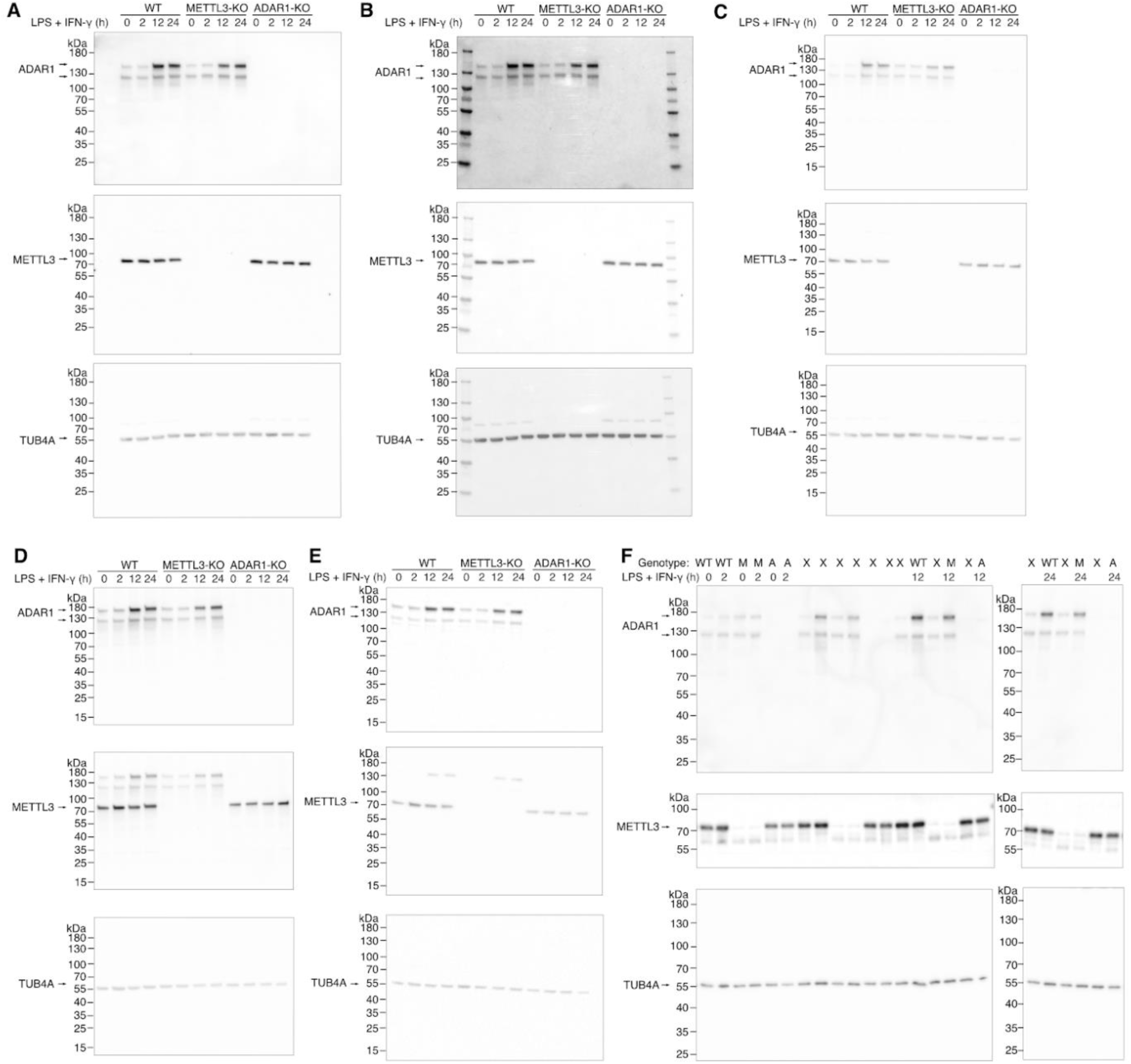
Western blots showing protein expression during 24 h time course upon LPS and IFN-γ treatment. (A-F) WT (wild type), METTL3-KO and ADAR1-KO macrophages were treated with LPS and IFN-γ. Protein samples were obtained at timepoint 0 h, 2 h, 12 h, and 24 h and analyzed by Western blot. Molecular (Mol.) size markers were included on each membrane, sizes as indicated. **(A, C-F)** Backgound-substracted membranes showing protein bands used for quantification. TUB4A served as loading control and was measured on the same membrane as proteins of interest. Each panel depicts an independent experiment. **(B)** Exemplary overlay of protein bands and molecular size markers presented in **(A). (F)** The two membranes were treated in parallel, samples were obtained from the same experiment and are labelled as followed: WT, wild type; M, METTL3-KO; A, ADAR1-KO; X, additional experimental conditions not relevant for the presented data.

**Suppl. Fig. 3.**
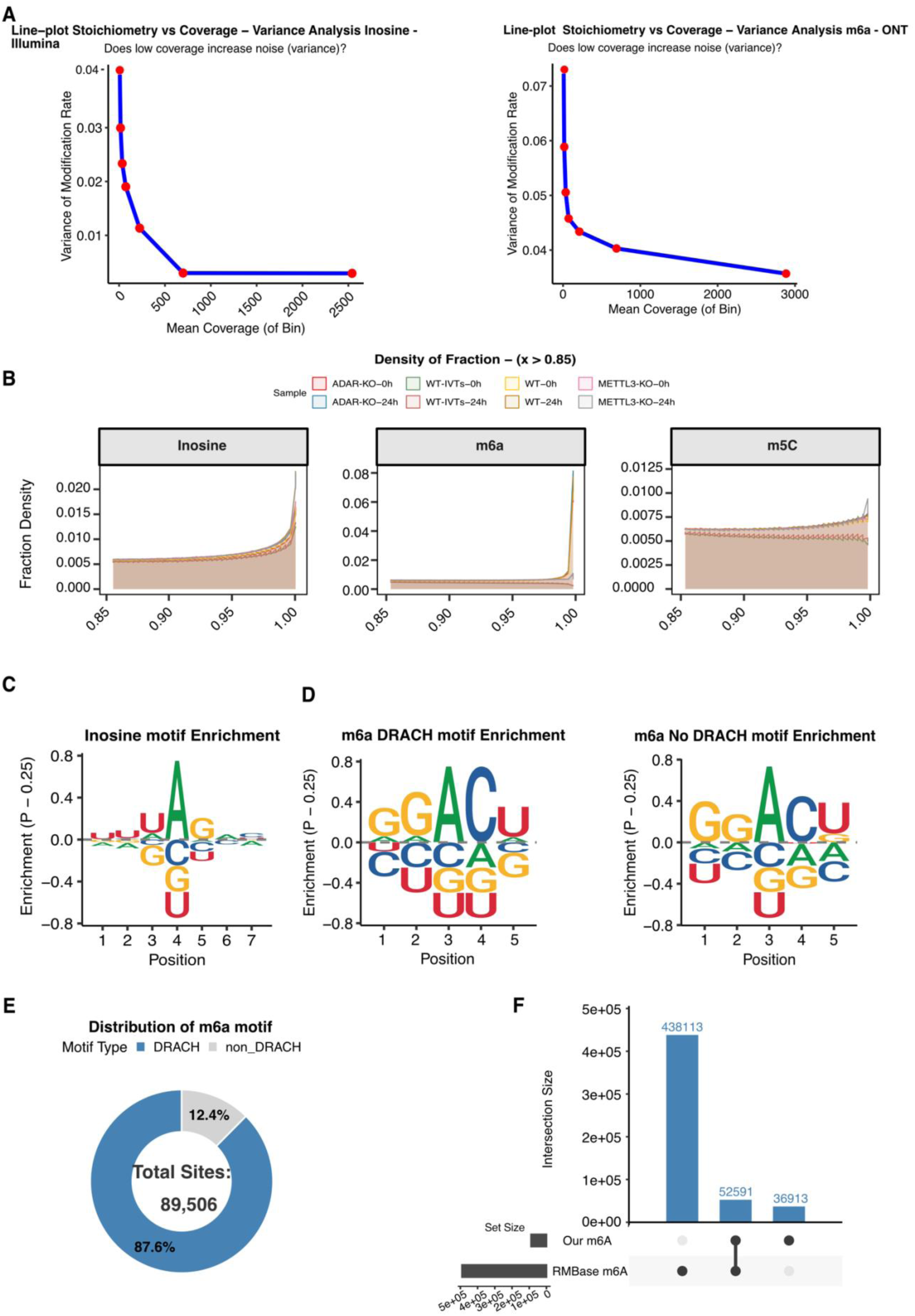
Nanopore Sequencing and Direct RNA Modification Metrology. (A) Stoichiometry Variance as a Function of Depth. Line plots illustrating the relationship between sequencing coverage depth (x-axis, Log_10_ scale) and the variance of stoichiometry for (left) detected A-to-I editing sites and (right) detected m6A modifications. For comparative analysis of stoichiometry between genotypes, sites were filtered by a minimum median coverage of 30 reads, a median of ≥5 modified reads, chosen to mitigate stoichiometry variance, and consistent detection across replicates and genotypes. This threshold partially mitigates the high variance in stoichiometry at low-coverage sites; although a coverage of 30 reads does not correspond to the point at which variance fully stabilizes, it was chosen as the highest threshold that avoids a substantial loss of data, thus balancing detection sensitivity with measurement stability across comparisons. Sites meeting these criteria are shown in blue. **(B) Distribution of base modification probabilities.** Density of modification calls as a function of the probability assigned by Modkit (sample-probs; x-axis, restricted to probability > 0.85) for inosine, m6A and m5C (separate panels) in each sample (colours). The y-axis (Fraction Density) shows the fraction of modification calls at a given probability. The x-axis is limited to the high-probability range because this is where differences between samples emerge. For m6A, WT samples separate from METTL3-KO samples, and for m5C, WT samples separate from the IVT control, supporting that high-probability calls reflect the presence of the modification. For inosine, no such separation is observed between genotypes, so Nanopore probabilities alone do not discriminate ADAR1-dependent editing; inosine sites were therefore additionally supported by Illumina-based editing calls (see Methods). IVT, in vitro transcribed (unmodified) control. **(C-D)** A-to-I and m6A Sequence Motifs. Sequence logos displaying the flanking nucleotide context (± 3 bp for A-to-I and ± 2 for m6A) around the modified base (0) for A-to-I editing sites and m6A sites. For m6A, the analysis confirms strong enrichment within the canonical **DRACH** motif (D=A/G/U, R=A/G, A=m6A, C, H=A/C/U). A detailed non-DRACH motif is also provided, illustrating the context for the minor subset of modifications. **(E) Motif Distribution of m6A Sites.** Donut chart showing the quantification of all detected m6A sites falling within the canonical **DRACH** motif (blue) vs. **Non-DRACH** motifs (light grey). Consistent with established epitranscriptomic biology, the vast majority (∼88%) of modifications align with the canonical DRACH sequence context. **(F) Benchmarking with RMBase v2.0.** Bar chart illustrating the intersection of the detected high-confidence m6A dataset with modifications cataloged in RMBase v2.0 (a database primarily of antibody-based (MeRIP-seq) peaks). The analysis shows a significant overlap of ∼60%, while ∼40% represent modifications detected exclusively in this study. While this exclusive subset likely reflects modifications within clusters or structured regions often inaccessible to antibody-based methods, highlighting the complementarity of antibody-independent Nanopore sequencing, it may also include a fraction of false positives inherent to de novo modification calling.

**Suppl. Fig. 4.**
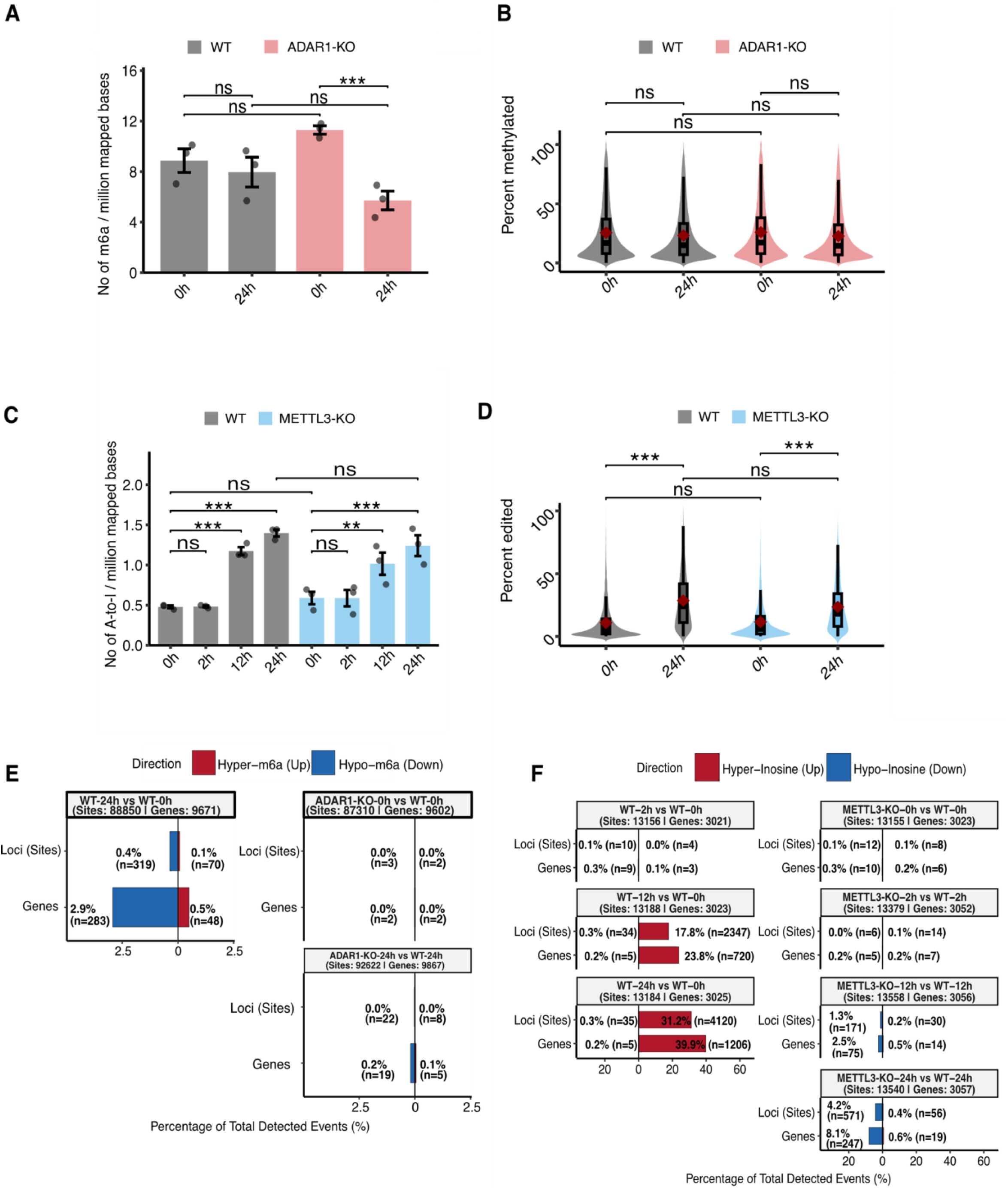
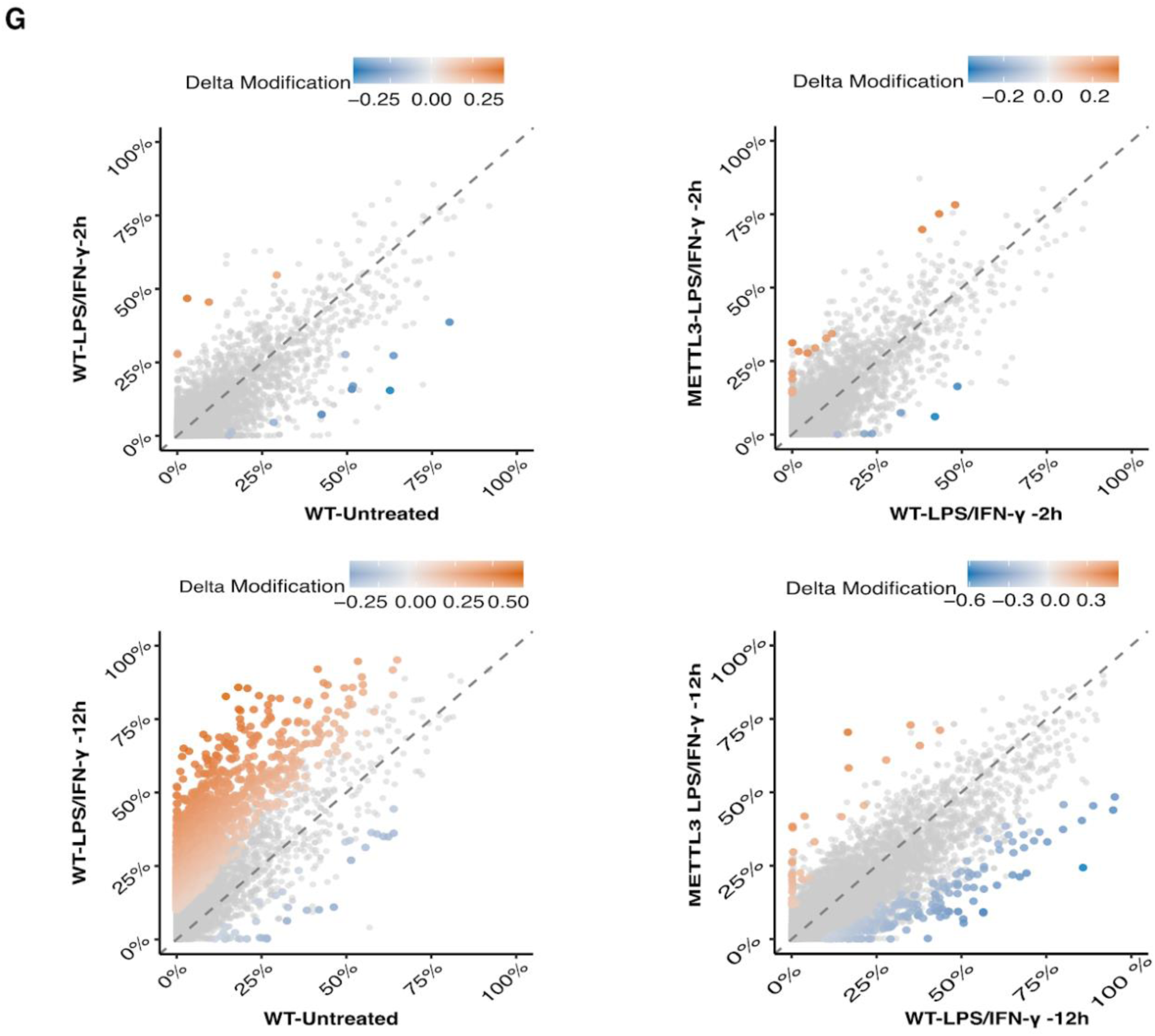
Global Dynamics and Distribution of m6A and A-to-I Editing. (A,C) Quantification of Global Modification Burdens. Barplots display the normalized counts of m6A **(A)** sites and A-to-I **(C)** editing sites across genotypes and time points. Modification counts are expressed as the number of sites per million mapped bases to enable reliable comparison across samples. Data represent the mean ± SEM of biological replicates. Statistical significance was assessed using a Generalized Linear Model (GLM) with a Quasi_Poisson distribution and a log-link function, accounting for the discrete nature of count data and biological overdispersion. Brackets indicate specific longitudinal comparisons within genotypes or between genotypes at matched time points. Significance levels: *p < 0.05, **p < 0.01, ***p < 0.001; ns, non-significant. **(B,D) Distribution of Modification Stoichiometry.** Violin plots with overlaid boxplots illustrate the global distribution of m6A **(B)** and A-to-I **(D)** editing stoichiometry. For this analysis, sites were strictly filtered to those with a minimum median sequencing coverage of 30 reads to mitigate stoichiometry variance, ≥5 modified reads, and replicates/genotypes consistency. Statistical significance was assessed using a Beta regression model within a Generalized Additive Modeling (GAM) framework, which is optimized for continuous proportional data bounded between 0 and 1. Random effects were included to account for biological variation across replicates. Significance levels: *p < 0.05, ***p < 0.001; ns, non-significant. **(E-F) Diverging Summaries of Differential Modifications.** Horizontal diverging bar plots represent the landscape of significantly altered m6A **(E)** and A-to-I editing **(F)** events. Data are partitioned into individual modification sites (Loci) and affected genes. Bar lengths indicate the percentage of significantly hyper-modified (red) and hypo-modified (blue) features relative to the total detected features. Significance for m6A and A-to-I editing was defined as required FDR < 0.05 and a stoichiometry change Δ Stoichiometry >= 10 %. **(G) Absolute Stoichiometry Comparisons of A-to-I Editing.** Scatterplots illustrating mean stoichiometry for individual A-to-I editing sites between specific experimental conditions: WT LPS-IFN-γ 2h vs. Untreated (top left), METTL3-KO LPS-IFN-γ 2h vs. WT LPS-IFN-γ 2h (top right), WT LPS-IFN-γ 12h vs. Untreated (bottom left) and METTL3-KO LPS-IFN-γ 12h vs. WT LPS-IFN-γ 12h (bottom right). Sites with significant changes (FDR < 0.05 and |Δ| ≥ 10%) are highlighted in color (orange: increased; blue: decreased), while stable sites are shown in grey. The diagonal identity line indicates no change in editing levels. **(A-G)** Data are derived from n=3 biological replicates per genotype and timepoint combination.

**Suppl. Fig. 5.**
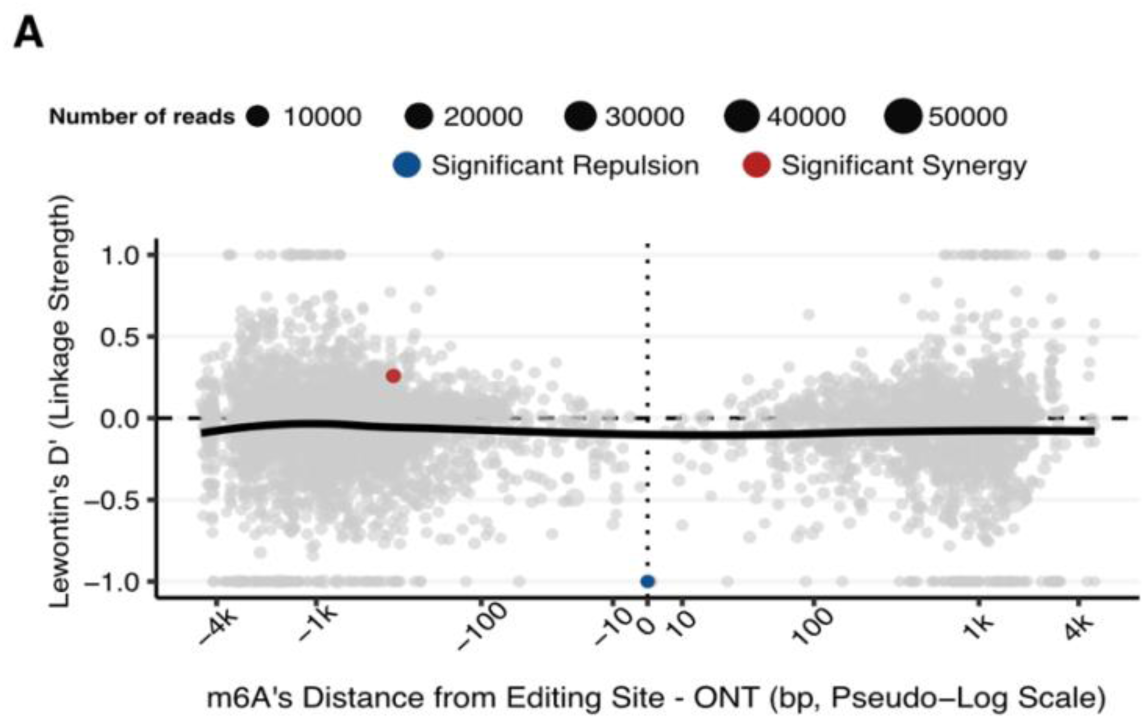
Co-occurrence of m6A and A-to-I Editing. (A) Linkage and co-occurrence of RNA modifications on single RNA molecules. (Full Dataset). Plot showing Lewontin’s D′ for all proximal modification pairs resolved on individual reads in the nanopore direct RNA sequencing dataset (844 inosine and 1,513 m6A sites across 181 genes). D′ was computed from reads spanning both sites of a pair (per-read modification probability ≥ 0.6; ≥ 30 spanning reads), where D′ = 0 denotes independence. Red points indicate significant co-occurrence on the same molecule (synergy, D′ > 0) and blue points significant mutual exclusivity (repulsion, D′ < 0); significance was called at FDR < 0.05 (Fisher’s exact test) with |D′| > 0.2. Grey points are pairs not reaching these thresholds. ONT, Oxford nanopore technology; LOESS, locally estimated scatterplot smoothing.

## References

1. Shi, H., Chai, P., Jia, R. & Fan, X. Novel insight into the regulatory roles of diverse RNA modifications: Re-defining the bridge between transcription and translation. Mol. Cancer 19, 78 (2020).

2. Meyer, K. D. et al. Comprehensive Analysis of mRNA Methylation Reveals Enrichment in 3′ UTRs and near Stop Codons. Cell 149, 1635–1646 (2012).

3. Dominissini, D. et al. Topology of the human and mouse m6A RNA methylomes revealed by m6A-seq. Nature 485, 201–206 (2012).

4. Liu, J. et al. A METTL3–METTL14 complex mediates mammalian nuclear RNA N6-adenosine methylation. Nat. Chem. Biol. 10, 93–95 (2014).

5. Jiang, X. et al. The role of m6A modification in the biological functions and diseases. Signal Transduct. Target. Ther. 6, 74 (2021).

6. Ke, S. et al. A majority of m6A residues are in the last exons, allowing the potential for 3’ UTR regulation. Genes Dev. 29, 2037–2053 (2015).

7. Zheng, G. et al. ALKBH5 is a mammalian RNA demethylase that impacts RNA metabolism and mouse fertility. Mol. Cell 49, 18–29 (2013).

8. Jia, G. et al. N6-Methyladenosine in nuclear RNA is a major substrate of the obesity-associated FTO. Nat. Chem. Biol. 7, 885–887 (2011).

9. Wang, X. et al. N6-methyladenosine Modulates Messenger RNA Translation Efficiency. Cell 161, 1388– 1399 (2015).

10. Wang, X. et al. N6-methyladenosine-dependent regulation of messenger RNA stability. Nature 505, 117– 120 (2014).

11. Meyer, K. D. et al. 5′ UTR m6A Promotes Cap-Independent Translation. Cell 163, 999–1010 (2015).

12. Zhou, J. et al. Dynamic m6A mRNA methylation directs translational control of heat shock response. Nature 526, 591–594 (2015).

13. Wang, H., Tang, A., Cui, Y., Gong, H. & Li, H. LRPPRC facilitates tumor progression and immune evasion through upregulation of m6A modification of PD-L1 mRNA in hepatocellular carcinoma. Front. Immunol. 14, 1144774 (2023).

14. Huang, H. et al. Recognition of RNA N6-methyladenosine by IGF2BP Proteins Enhances mRNA Stability and Translation. Nat. Cell Biol. 20, 285–295 (2018).

15. Alarcón, C. R. et al. HNRNPA2B1 is a mediator of m6A-dependent nuclear RNA processing events. Cell 162, 1299–1308 (2015).

16. Roundtree, I. A. et al. YTHDC1 mediates nuclear export of N6-methyladenosine methylated mRNAs. eLife 6, e31311 (2017).

17. Gao, Y. et al. m6A Modification Prevents Formation of Endogenous Double-Stranded RNAs and Deleterious Innate Immune Responses during Hematopoietic Development. Immunity 52, 1007–1021.e8 (2020).

18. Qiu, W. et al. N6-methyladenosine RNA modification suppresses antiviral innate sensing pathways via reshaping double-stranded RNA. Nat. Commun. 12, 1582 (2021).

19. Bass, B. L. RNA Editing by Adenosine Deaminases That Act on RNA. Annu. Rev. Biochem. 71, 817–846 (2002).

20. Wulff, B.-E. & Nishikura, K. Substitutional A-to-I RNA editing. Wiley Interdiscip. Rev. RNA 1, 90–101 (2010).

21. Basilio, C., Wahba, A. J., Lengyel, P., Speyer, J. F. & Ochoa, S. SYNTHETIC POLYNUCLEOTIDES AND THE AMINO ACID CODE, V*. Proc. Natl. Acad. Sci. U. S. A. 48, 613–616 (1962).

22. Patterson, J. B. & Samuel, C. E. Expression and regulation by interferon of a double-stranded-RNA-specific adenosine deaminase from human cells: evidence for two forms of the deaminase. Mol. Cell. Biol. 15, 5376–5388 (1995).

23. George, C. X., John, L. & Samuel, C. E. An RNA Editor, Adenosine Deaminase Acting on Double-Stranded RNA (ADAR1). J. Interferon Cytokine Res. 34, 437–446 (2014).

24. Mannion, N. M. et al. The RNA-Editing Enzyme ADAR1 Controls Innate Immune Responses to RNA. Cell Rep. 9, 1482–1494 (2014).

25. Pestal, K. et al. Isoforms of RNA-Editing Enzyme ADAR1 Independently Control Nucleic Acid Sensor MDA5-Driven Autoimmunity and Multi-organ Development. Immunity 43, 933–944 (2015).

26. Wang, J., Yan, S., Lu, H., Wang, S. & Xu, D. METTL3 Attenuates LPS-Induced Inflammatory Response in Macrophages via NF-κB Signaling Pathway. Mediators Inflamm. 2019, 3120391 (2019).

27. Zong, X. et al. Mettl3 Deficiency Sustains Long-Chain Fatty Acid Absorption through Suppressing Traf6-Dependent Inflammation Response. J. Immunol. Author Choice 202, 567–578 (2019).

28. Wang, X. et al. The m6A Reader IGF2BP2 Regulates Macrophage Phenotypic Activation and Inflammatory Diseases by Stabilizing TSC1 and PPARγ. Adv. Sci. 8, 2100209 (2021).

29. Cai, D., Sun, C., Murashita, T., Que, X. & Chen, S.-Y. ADAR1 non-editing function in macrophage activation and abdominal aortic aneurysm. Circ. Res. 132, e78–e93 (2023).

30. Sun, C., Cai, D. & Chen, S.-Y. ADAR1 promotes systemic sclerosis via modulating classic macrophage activation. Front. Immunol. 13, 1051254 (2022).

31. Liu, Y. et al. The N6-methyladenosine (m6A)-forming enzyme METTL3 facilitates M1 macrophage polarization through the methylation of STAT1 mRNA. Am. J. Physiol. Cell Physiol. 317, C762–C775 (2019).

32. Schott, J. et al. Translational Regulation of Specific mRNAs Controls Feedback Inhibition and Survival during Macrophage Activation. PLOS Genet. 10, e1004368 (2014).

33. Medzhitov, R. & Horng, T. Transcriptional control of the inflammatory response. Nat. Rev. Immunol. 9, 692–703 (2009).

34. Poh, H. X., Mirza, A. H., Pickering, B. F. & Jaffrey, S. R. Alternative splicing of METTL3 explains apparently METTL3-independent m6A modifications in mRNA. PLOS Biol. 20, e3001683 (2022).

35. Pecori, R. et al. ADAR1-mediated RNA editing promotes B cell lymphomagenesis. iScience 26, 106864 (2023).

36. Roche, P. A. & Furuta, K. The ins and outs of MHC class II-mediated antigen processing and presentation. Nat. Rev. Immunol. 15, 203–216 (2015).

37. Brown, G. D. & Gordon, S. A new receptor for β-glucans. Nature 413, 36–37 (2001).

38. Xuan, J.-J. et al. RMBase v2.0: deciphering the map of RNA modifications from epitranscriptome sequencing data. Nucleic Acids Res. 46, D327–D334 (2018).

39. Lewontin, R. C. The Interaction of Selection and Linkage. I. General Considerations; Heterotic Models. Genetics 49, 49–67 (1964).

40. Acera Mateos, P., et al. Prediction of m6A and m5C at single-molecule resolution reveals a transcriptome-wide co-occurrence of RNA modifications. Nat. Commun. 15, 3899 (2024).

41. Byrne, A. et al. Direct RNA Sequencing reveals epitranscriptomic regulation of brain cells and Alzheimer’s Disease pathology. 2026.05.18.724443 Preprint at 10.64898/2026.05.18.724443 (2026).

42. Orji, O. C. et al. Global Co-regulatory Cross Talk Between m6A and m5C RNA Methylation Systems Coordinate Cellular Responses and Brain Disease Pathways. Mol. Neurobiol. 62, 5006–5021 (2025).

43. Tassinari, V. et al. ADAR1 is a new target of METTL3 and plays a pro-oncogenic role in glioblastoma by an editing-independent mechanism. Genome Biol. 22, 51 (2021).

44. Terajima, H., et al. N6-methyladenosine promotes induction of ADAR1-mediated A-to-I RNA editing to suppress aberrant antiviral innate immune responses. PLOS Biol. 19, e3001292 (2021).

45. Xiao, W. et al. Nuclear m(6)A Reader YTHDC1 Regulates mRNA Splicing. Mol. Cell 61, 507–519 (2016).

46. Rabinovici, R. et al. ADAR1 Is Involved in the Development of Microvascular Lung Injury. Circ. Res. 88, 1066–1071 (2001).

47. Yang, J.-H. et al. Widespread inosine-containing mRNA in lymphocytes regulated by ADAR1 in response to inflammation. Immunology 109, 15–23 (2003).

48. Li, Y., et al. RNA Editing Enzyme ADAR1 Regulates METTL3 in an Editing Dependent Manner to Promote Breast Cancer Progression via METTL3/ARHGAP5/YTHDF1 Axis. Int. J. Mol. Sci. 23, 9656 (2022).

49. Nguyen, T. A. et al. Direct identification of A-to-I editing sites with nanopore native RNA sequencing. Nat. Methods 19, 833–844 (2022).

50. Kellner, S. et al. Absolute and relative quantification of RNA modifications via biosynthetic isotopomers. Nucleic Acids Res. 42, e142 (2014).

51. Hengesbach, M. et al. Toward standardized epitranscriptome analytics: an inter-laboratory comparison of mass spectrometric detection and quantification of modified ribonucleosides in human RNA. Nucleic Acids Res. 53, gkaf895 (2025).

52. Love, M. I., Huber, W. & Anders, S. Moderated estimation of fold change and dispersion for RNA-seq data with DESeq2. Genome Biol. 15, 550 (2014).

53. Bolger, A. M., Lohse, M. & Usadel, B. Trimmomatic: A flexible trimmer for Illumina sequence data. Bioinformatics 30, 2114–2120 (2014).

54. Dobin, A. et al. STAR: Ultrafast universal RNA-seq aligner. Bioinformatics 29, 15–21 (2013).

55. Okonechnikov, K., Conesa, A. & García-Alcalde, F. Qualimap 2: Advanced multi-sample quality control for high-throughput sequencing data. Bioinformatics 32, 292–294 (2016).

56. Li, H. et al. The Sequence Alignment/Map format and SAMtools. Bioinformatics 25, 2078–2079 (2009).

57. Flati, T. et al. HPC-REDItools: a novel HPC-aware tool for improved large scale RNA-editing analysis. BMC Bioinformatics 21, 353 (2020).

58. Picardi, E. & Pesole, G. REDItools: High-throughput RNA editing detection made easy. Bioinformatics 29, 1813–1814 (2013).

59. Lawrence, M. et al. Software for Computing and Annotating Genomic Ranges. PLOS Comput. Biol. 9, e1003118 (2013).

60. Wu, H., Wang, C. & Wu, Z. A new shrinkage estimator for dispersion improves differential expression detection in RNA-seq data. Biostatistics 14, 232–243 (2013).

61. Park, Y. & Wu, H. Differential methylation analysis for BS-seq data under general experimental design. Bioinformatics 32, 1446–1453 (2016).

62. Feng, H., Conneely, K. N. & Wu, H. A Bayesian hierarchical model to detect differentially methylated loci from single nucleotide resolution sequencing data. Nucleic Acids Res. 42, e69 (2014).

63. Patro, R., Duggal, G., Love, M. I., Irizarry, R. A. & Kingsford, C. Salmon provides fast and bias-aware quantification of transcript expression. Nat. Methods 14, 417–419 (2017).

64. Robinson, M. D., McCarthy, D. J. & Smyth, G. K. edgeR: a Bioconductor package for differential expression analysis of digital gene expression data. Bioinforma. Oxf. Engl. 26, 139–40 (2010).

65. Soneson, C., Love, M. I. & Robinson, M. D. Differential analyses for RNA-seq: transcript-level estimates improve gene-level inferences. Preprint at 10.12688/f1000research.7563.1 (2016).

66. Lewis M. B. K., Rana S. EnhancedVolcano: Publication-ready volcano plots with enhanced colouring and labeling. https://github.com/kevinblighe/EnhancedVolcano. (2024).

67. Wu, T. et al. clusterProfiler 4.0: A universal enrichment tool for interpreting omics data. The Innovation 2, 100141 (2021).

68. Li, H. Minimap2: pairwise alignment for nucleotide sequences. Bioinformatics 34, 3094–3100 (2018).

69. McLaren, W. et al. The Ensembl Variant Effect Predictor. Genome Biol. 17, 122 (2016).

70. Zhang, F. & Chen, J. Y. HOMER: a human organ-specific molecular electronic repository. BMC Bioinformatics 12, S4 (2011).

71. Chen, N. Using RepeatMasker to identify repetitive elements in genomic sequences. Curr. Protoc. Bioinforma. Chapter 4, Unit 4.10 (2004).

72. Wood, S. N. Fast stable restricted maximum likelihood and marginal likelihood estimation of semiparametric generalized linear models. J. R. Stat. Soc. Ser. B Stat. Methodol. 73, 3–36 (2011).

73. Searle, S. R., Speed, F. M. & Milliken, G. A. Population Marginal Means in the Linear Model: An Alternative to Least Squares Means. Am. Stat. 34, 216–221 (1980).

